# Peptide-Based Complex Coacervates Stabilized by Cation-π Interactions for Cell Engineering

**DOI:** 10.1101/2024.10.20.619264

**Authors:** Yue Sun, Xi Wu, Jianguo Li, Chandra Shekhar Verma, Jing Yu, Ali Miserez

## Abstract

Complex coacervation is a form of liquid-liquid phase separation (LLPS), whereby two types of macromolecules, usually bearing opposite net charges, self-assemble into dense microdroplets driven by weak molecular interactions. Peptide-based coacervates have recently emerged as promising carriers to deliver large macromolecules (nucleic acids, proteins and complex thereof) inside cells. Thus, it is essential to understand their assembly/disassembly mechanisms at the molecular level in order to tune the thermodynamics of coacervates formation and the kinetics of cargo release upon entering the cell. In this study, we design histidine-rich peptides consisting of modular sequences in which we systematically incorporate cationic, anionic, or aromatic residues at specific positions along the sequence in order to modulate intermolecular interactions and the resulting coacervation stability. We show that cation-π interactions between arginine and aromatic side chains are particularly efficient in stabilizing complex coacervates, and these interactions can be disrupted in the protein-rich intracellular environment, triggering the disassembly of complex coacervates followed by cargo release. With the additional grafting of a disulfide-based self-immolative sidechain, these complex coacervates exhibit enhanced stability and can deliver proteins, mRNA, and CRISPR/Cas9 genome editing tools with tunable release kinetics into cells. This capability extends to challenging cell types, such as macrophages. Our study highlights the critical role of cation-π interactions in the design of peptide-based coacervates, expanding the biomedical and biotechnology potential of this emerging innovative intracellular delivery platform.

**Table of Content:** Tandem peptides designed from histidine-rich beak proteins are modified with either negative, positive, or aromatic residues that can undergo complex coacervation through cation-_π_ rather than electrostatic interactions. These complex coacervates can effectively recruit various macromolecular cargos and deliver them intracellularly with controlled release kinetics, including in hard-to-transfect macrophages.

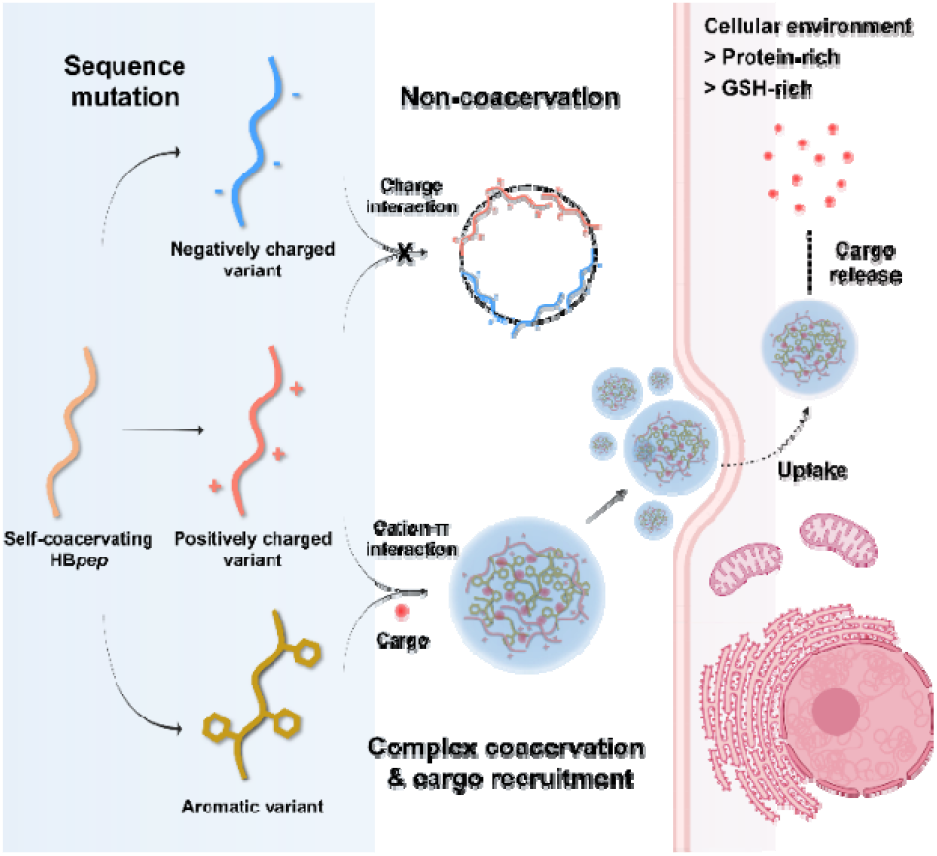

## Introduction

Weak inter- and intra-molecular interactions are fundamental to self-assembly processes that drive the formation of complex biological structures.^[1-5]^ Among these, coacervation is a liquid-liquid phase separation (LLPS) process resulting in the formation of macromolecular-rich microdroplets whose assembly is governed by diverse weak interactions.^[6-9]^ Coacervation can be classified into self-coacervation and complex coacervation, each with distinct mechanisms and molecular interaction profiles.^[10-11]^ Self-coacervation involves interactions between homotypic macromolecules, where hydrophobic, ionic, and hydrogen bonding interactions drive phase separation, triggered by a change in the local microenvironment such as pH, temperature, or ionic strength.^[12-13]^ For example, the self-coacervation of elastin-like polypeptides (ELPs) arises from hydrophobic interactions that are favored above a critical temperature.^[14-16]^ In contrast, complex coacervation involves interactions between heterotypic macromolecules, typically oppositely charged polyelectrolytes (*e*.*g*. proteins, nucleic acids, or synthetic polymers), leading to phase separation by complementary ionic interactions.^[17-19]^ However, these interactions are often more complex, involving a combination of electrostatic forces, hydrophobic effects, and specific non-covalent interactions such as cation-π interactions.^[20-21]^

Typical examples of self-coacervation are histidine-rich beak proteins (HBP), which our team has studied in recent years, identifying the relationships between their sequence and coacervation mechanisms.^[22-25]^ The modular and low sequence complexity nature of HBP makes it an excellent model system to explore LLPS and underlying molecular mechanisms. Inspired by these studies, we have designed a self-coacervating peptide, HB*pep*, composed of five repeats of Gly-His-Gly-**X**-Tyr (GHGXY) and a C-terminal Trp (W) to enhance and stabilize coacervate formation (Table S1). In this sequence, the X position is a variable residue and occupied by Val (V), Pro (P), or Leu (L) in the natural HBP sequence. This design provides an ideal framework for investigating the role of specific residue substitutions on the coacervation behavior of peptide variants.^[26-27]^

Beyond its structural versatility, HB*pep* has been demonstrated to be highly efficient at forming coacervates and recruit a wide range of cargos including proteins, peptides, chemotherapy drugs, and inorganic nanoparticles.^[28-29]^ To expand its functionality, we introduced a redox-responsive release mechanism by incorporating a positively charged lysine (Lys, K) residue that disrupts self-coacervation under physiological conditions. Further functionalization of the added Lys with a disulfide-based self-immolative moiety resulted in the peptide HB*pep*-K^SP^, which can encapsulate and deliver therapeutic cargos intracellularly, with release triggered by endogenous glutathione (GSH) (Figure S1).^[30-31]^ While this delivery platform has the potential to deliver modalities across a broad range of therapeutic applications, it still faces significant challenges. This includes the slow release kinetics attributed to the hydrophobic nature of the peptide sequence, which may hinder water and GSH accessibility, thus affecting the efficiency and kinetics of cargo release. However, hydrophobic interactions are crucial for the self-coacervation process,^[24, 32]^ highlighting the challenge of balancing stability and subsequent disassembly of the coacervates, as well as cargo release. Thus, complex coacervates with more hydrophilic interior could offer an improved solution, facilitating the rapid release of cargos while maintaining coacervate integrity.

In this study, we introduced cationic (Lys and Arg; K and R), anionic (Glu; E), and aromatic (Tyr; Y) residues at the X position of HB*pep* to assess their influence on self- and complex coacervation of the resulting peptide variants (Figure 1a). Our findings reveal that variants containing cationic (K, R) and anionic (E) residues tend to remain in solution across all tested pHs and peptide concentrations, while aromatic (Y) residues promote aggregation. Furthermore, mixing peptide variants with oppositely charged residues at the X position was insufficient to induce complex coacervation through electrostatic interactions. However, cation-π interactions induced by mixing Arg- and Tyr-containing variants were found to promote phase separation and the formation of complex coacervates. Notably, these cation-π interactions can be disrupted by the high protein content within the crowded intracellular environment, providing an additional strategy for intracellular release of macromolecular cargos.

**Figure 1.**
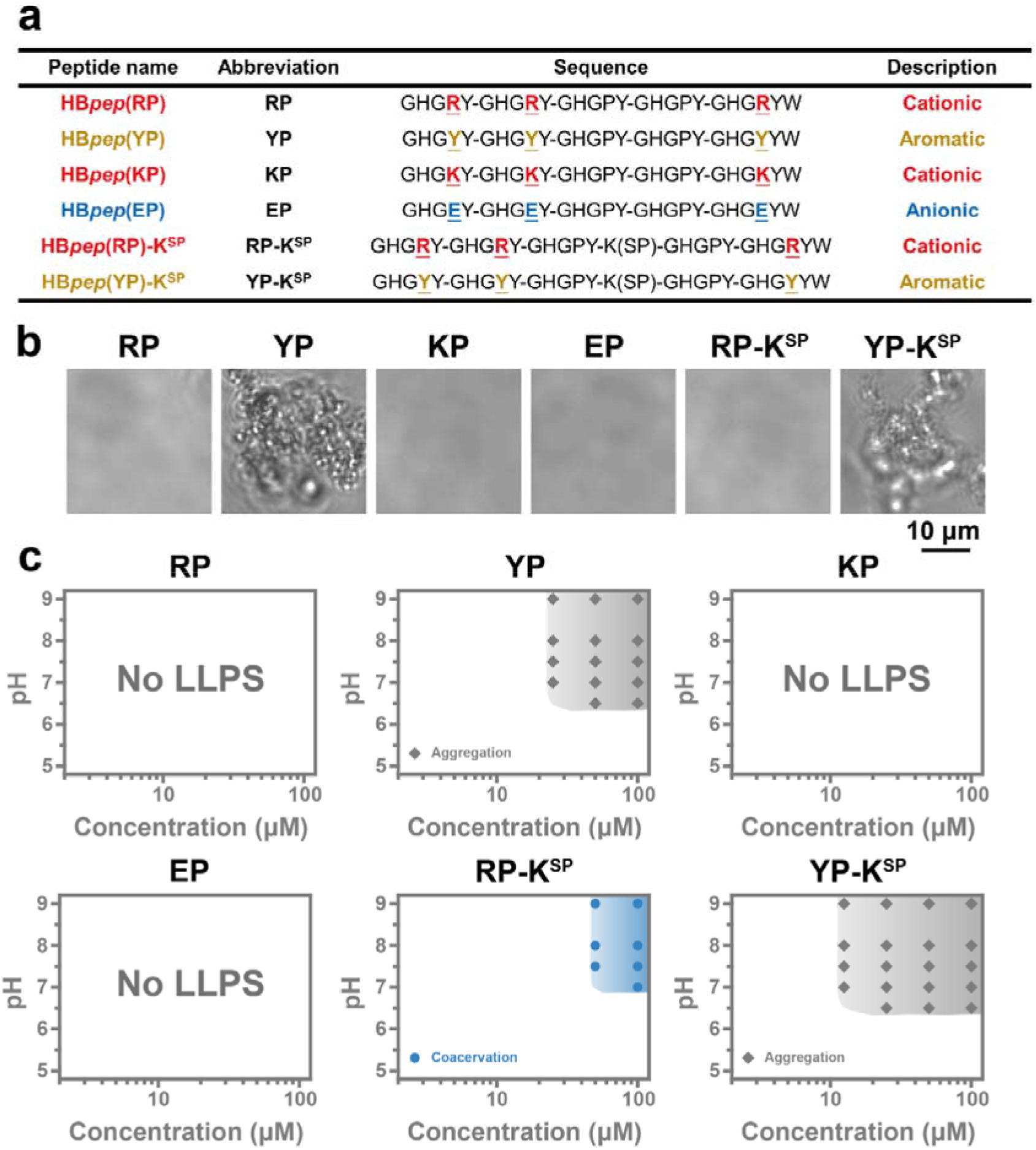
Sequence and self-coacervation of HB*pep* variants and derivates. **(a)** Peptide sequences of HB*pep* and HB*pep*-K^SP^ derivates modified by a self-immolative moiety on the lysine (K) side chain. The abbreviations refer to the X positions in the pentapeptide repeat GHGXY. **(b)** Representative optical micrographs of variants (50 μM, pH = 7.0, IS = 100 mM). **(c)** Phase diagram of variants at the ionic strength (IS) of 100 mM, with the region of coacervation or aggregation shadowed in blue and grey, respectively.

Furthermore, by incorporating a disulfide bond and self-immolative moiety into the peptide variants, we engineered complex coacervates capable of rapidly releasing functional macromolecules under the reducing and crowded intracellular environment. Adjusting the ratio of cation/π-rich variants in the complex coacervates allowed us to fine-tune cargo release kinetics and delivery efficiencies. These complex coacervates demonstrated robust delivery efficiency for a wide range of macromolecular therapeutics, including proteins, antibodies, mRNA, and CRISPR/Cas9 genome editing tools, even in macrophages that are typically highly resistant to transfection. Using this capability, we seamlessly delivered CRISPR/Cas9 editing tools to knock down the signal-regulatory protein alpha (SIRPα) of macrophages, which inhibits phagocytosis of tumor cells.^[33-34]^ This study shows that in addition to their self-assembly into simple coacervates, the HB*pep* family can be expanded to form complex coacervates. By balancing the weak non-bonded cation-π interactions, the formation and stability of these coacervates can be modulated to precisely control cargo release, broadening their scope of applications in chemical biology, drug delivery, and cell therapy.

## Results and Discussion

### Preparation and Coacervation Study of Peptide Variants Derived from HB*pep*

To investigate the effect of residue substitutions on the coacervation of phase-separating peptides, four variants were synthesized by mutating the 4th, 9th, and 24th positions of the HB*pep* sequence (Figure 1a). Three variants containing charged residues, namely RP, KP, and EP, did not undergo phase separation at concentrations up to 100 μM across a broad pH range of 5 to 9 (Figures 1b-c). Conversely, the Y-containing variant YP, tended to form aggregates due to strong hydrophobic interactions and π-π stacking (Figures 1b-c).

Additionally, two derivative peptides incorporating Lys at the 16th position followed by grafting it with the self-immolative modification were synthesized from the RP and YP backbones (Figures 1a and S1). These derivatives, termed RP-K^SP^ and YP-K^SP^, displayed phase separation behaviors similar to their parent peptides, RP and YP. However, due to the increased hydrophobicity of the modification, RP-K^SP^ formed coacervates at higher concentrations and pH values, whereas YP-K^SP^ aggregated at lower concentrations compared to YP (Figures 1b-c).

Building on these results, we assessed the complex coacervation behavior of peptide variants. Unlike the favorable electrostatic interactions typically seen in classic poly-lysine and poly-glutamic acid systems,^[35-37]^ neither RP nor KP could engage in strong interactions with EP to drive the formation of complex coacervates (Figures 2a and S2a-b). A likely explanation is that the presence of only three charged residues per peptide is insufficient to establish stable inter-peptide interactions, while simultaneously weakening hydrophobic interactions. As a result, these variants not only failed to undergo self-coacervation but are also unable to form complex coacervates driven by electrostatic interactions.

**Figure 2.**
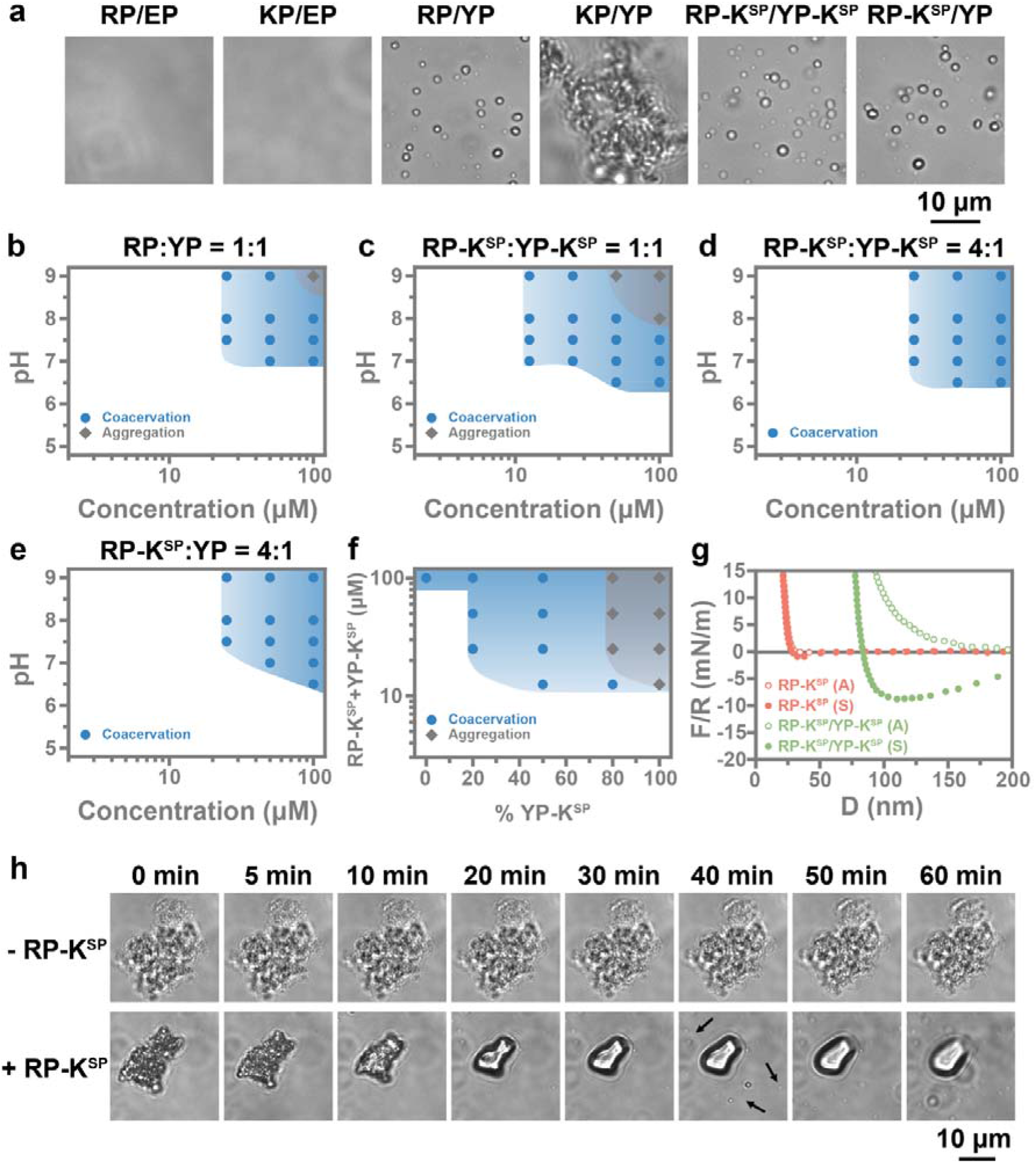
Complex coacervation of HB*pep* variants and derivates. **(a)** Representative optical micrographs of variant mixtures (total concentration = 50 μM, mix ratio = 1:1, pH = 7.0, IS = 100 mM). **(b-e)** Phase diagram of variant mixtures including RP:YP = 1:1 **(b)**, RP-K^SP^:YP-K^SP^ = 1:1 **(c)**, RP-K^SP^:YP-K^SP^ = 4:1 **(d)**, and RP-K^SP^:YP = 4:1 **(e)** at the ionic strength (IS) of 100 mM, with the region of coacervation or aggregation shadowed in blue and grey, respectively. **(f)** Phase diagram of RP-K^SP^:YP-K^SP^ mixed at different ratios. With increased YP-K^SP^ percentage, the coacervation region (blue-shadowed) expands and then becomes aggregation (grey-shadowed). **(g)** Normalized force-distance (*F/R* vs *D*) curves measured by SFA during the approach (A) and separate (S) of two cross-cylinder mica surfaces with RP-K^SP^ (50 μM in PBS) and RP-K^SP^:YP-K^SP^ = 1:1 (total concentration = 50 μM, mix ratio = 1:1, in PBS) in between. **(h)** Optical micrographs of aggregates formed by YP-K^SP^ evolve with/without RP-K^SP^ addition. The arrows indicate the formation of coacervates after adding the RP-K^SP^.

In contrast, RP and YP readily formed complex coacervates at a 1:1 molar ratio across a broad range of concentrations and pH values, as shown in Figures 2a-b. However, KP failed to prevent the aggregation of YP (Figures 2a and S2c), highlighting that the Lys-Tyr cation-π pair is insufficient to mitigate aggregation. In contrast, the cation-π interactions emerging from Arg-Tyr pairs is much stronger and can compete with π-π stacking between Tyr-Tyr pairs, resulting in intermolecular force balance that enables to form stable complex coacervates.^[38-40]^ Similar to their parent peptides, the RP-K^SP^ and YP-K^SP^ mixtures also formed coacervates under these conditions, but began to aggregate at pH 8 due to the hydrophobic nature of the K^SP^ moiety (Figures 2c). However, this aggregation could be further mitigated by increasing the proportion of RP-K^SP^ in the mixture (Figure 2d). Moreover, RP-K^SP^ and YP-K^SP^ can also interact with their less hydrophobic parent peptides, RP and YP, and influence the threshold concentration required to induce complex coacervation (Figures 2e and S2d). Based on this design, the phase behavior of the complexes could be fine-tuned by adjusting the cationic-to-aromatic peptide ratio, resulting in various phase behaviors ranging from no phase separation to complex coacervation and, ultimately, aggregation (Figure 2f). This ability to modulate the phase behavior highlights the versatility of this peptide family to form complex coacervates under varying conditions.

To further validate the formation of complex coacervates, we evaluated the viscoelastic characteristics of the RP-K^SP^/YP-K^SP^ complex using the surface force apparatus (SFA). The SFA employs two mica surfaces affixed on perpendicularly oriented cylindrical discs of radii *R*, where the coacervates form a liquid bridge in between. The cross-cylinders are approached with sub-nm distance resolution and then retracted, while the normal force between the cylinders is monitored with μN force sensitivity.^[41-42]^ Upon retraction, negative normalized forces (*F/R*) correspond to attractive interactions between the cross-cylinders. As shown in Figure 2g, RP-K^SP^ alone did not exhibit adhesive force, consistent with its weak self-coacervation ability. However, the RP-K^SP^/YP-K^SP^ complex demonstrated a strong adhesive force of - 8.78 mN/m, as well as clear hysteresis between approach and separation, both hallmark signatures of coacervates.^[43-44]^

Intriguingly, the cationic RP-K^SP^ not only prevented the aggregation of aromatic YP-K^SP^, but could also dissolve the pre-formed peptide aggregates. Figure 2h shows time-lapse microscopy images of YP-K^SP^ aggregates changing in size and texture, which evolved over time into coacervate microdroplets, whereas the aggregates remained intact in the absence of RP-K^SP^. Structural analyses using attenuated total reflection-Fourier transformed infrared spectroscopy (ATR-FTIR) revealed that YP-K^SP^ formed β-turn structures, characterized by the amide I peak centered at 1676 cm^-1^.^[45]^ In contrast, the amide I peak of the RP-K^SP^/YP-K^SP^ mixture at the molar ratio of 1:1 shifted to 1646 cm^-1^, indicative of disordered structures similar to homotypic RP-K^SP^ (Figure S3).^[46]^ These findings suggest strong interactions between the cationic and aromatic peptides, which facilitate complex coacervation and inhibit or even reverse aggregation. This is particularly stimulating, as misfolding of proteins into amyloid aggregates are central to many protein-related diseases.^[47-48]^ These results suggest a potential strategy to mitigate or reverse aggregation by harnessing cation-π interactions in aggregation-prone peptides.

### Cation-π Interactions Modulate Complex Coacervates Formation

To elucidate the role of weak non-bonded inter-molecular interactions between cationic and aromatic variants and derivatives in complex coacervation, we employed SFA measurements and molecular dynamics (MD) simulations. Symmetric SFA measurements –whereby the same peptide layer is coated on both cylinders– were conducted for RP-K^SP^ and YP-K^SP^ to quantitatively evaluate interactions between the same peptides. Notably, the adhesion force between YP-K^SP^ layers was approximately 1.4-fold higher compared to that between RP-K^SP^ layers (Figures 3a and S4). Moreover, the force-distance curves of YP-K^SP^ layers were not affected by the addition of tetramethylammonium hydroxide (TMA), an inhibitor of cation-π interactions, likely due to the absence of positively charged residues in its sequence. In contrast, the average adhesion force between RP-K^SP^ layers slightly decreased from 7.34 ± 0.10 mN/m to 6.67 ± 0.93 mN/m upon TMA addition, suggesting weak perturbation of cation-π interactions likely arising between Arg and Tyr of the GHGRY repeats as well as Arg and the C-terminal Trp (Figures 3a and S4). Asymmetric SFA measurements between RP-K^SP^ and YP-K^SP^ layers exhibited the strongest adhesion force of 11.69 ± 0.34 mN/m, which significantly decreased by 55% with the addition of 100 mM TMA (Figures 3a and S4). This result suggests that strong cation-π interactions occur between these peptide variants, which may prevent YP-K^SP^ from aggregating and instead favor the formation of YP-K^SP^/RP-K^SP^ complex coacervates.

**Figure 3.**
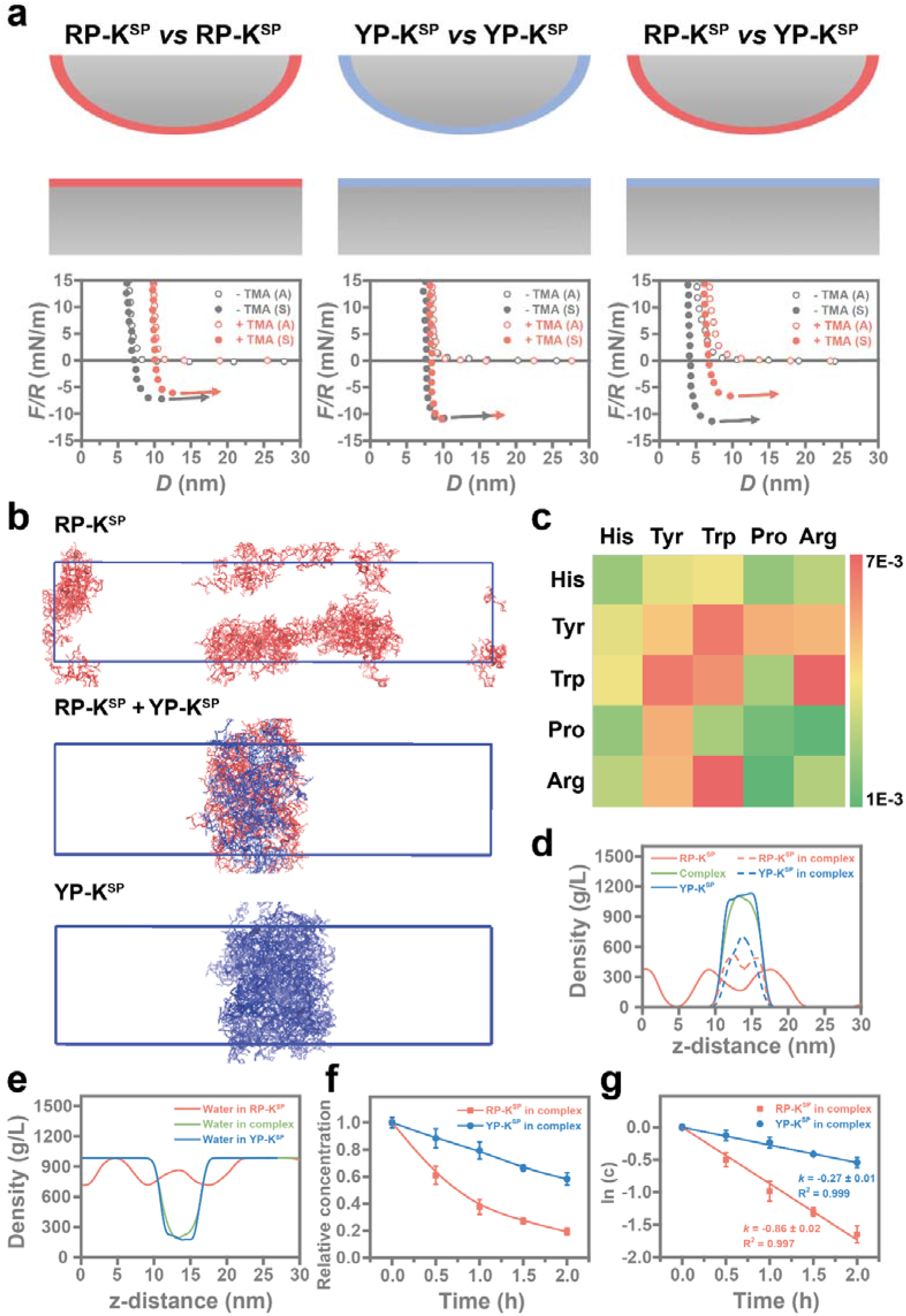
Molecular interactions in RP-K^SP^/YP-K^SP^ complex coacervation. **(a)** Normalized force-distance (*F/R* vs *D*) curves measured by SFA during the approach (A) and separation (S) of two cross-cylinder mica surfaces coated with RP-K^SP^ or YP-K^SP^ layers. Measurements were conducted in buffer at pH = 7.0 and IS = 100 mM with (red) or without (grey) the addition of 100 mM TMA. **(b)** Snapshots of the slab simulation of RP-K^SP^, RP-K^SP^/YP-K^SP^ mixture, and YP-K^SP^, indicate that the mixture and YP-K^SP^ form a single stable cluster, whereas RP-K^SP^ stays as isolated oligomers. **(c)** Quantification of contacts between different residues normalized by the number of residue-residue pairs in RP-K^SP^/YP-K^SP^ mixture systems. **(d-e)** The density of peptides **(d)** and water **(e)** in RP-K^SP^, RP-K^SP^/YP-K^SP^ mixture, and YP-K^SP^ clusters, analyzed from slab simulations. **(f)** Concentration decay of RP-K^SP^ and YP-K^SP^ peptide in the RP-K^SP^:YP-K^SP^ = 1:1 complex coacervates reduced by 1 mM GSH over time. **(g)** Concentration decay in natural logarithmic scale plotted as a function of time. The reaction rate constant *k* was obtained from the slopes of the fitted lines. Data are presented as the mean □±□ SD of *n* = 3 independent experiments.

MD simulations provided further insights into peptide assembly and peptide/peptide interactions at the molecular level. Simulations using both slab and cubic boxes indicated low propensity to form a compact cluster for RP-K^SP^ peptides, whereas YP-K^SP^ peptides and the 1:1 mixture of RP-K^SP^/YP-K^SP^ peptides rapidly collapsed into stable clusters (Figures 3b and S5). The residue contact map of the RP-K^SP^/YP-K^SP^ mixture and the total number of cation-π interactions confirmed strong cation-π interactions for the pairs Arg-Tyr and Arg-Trp (Figures 3c and S6a), consistent with the SFA measurements (Figure 3a).

From simulations using a slab box, the peptide density profile showed a single peak for both YP-K^SP^ and RP-K^SP^/YP-K^SP^ mixture, while multiple small peaks appeared for RP-K^SP^, corresponding to several small clusters (Figures 3d and 3e). Within the peptide coacervates, water density decreased but did not drop to zero, indicating the presence of significant numbers of “internal” water molecules. Further analysis of the RP-K^SP^/YP-K^SP^ mixture revealed distinct spatial distributions, with YP-K^SP^ preferentially located in the interior of the coacervates and RP-K^SP^ positioned at the coacervates-water interface (dashed line in Figure 3d). This distribution arises from differences in hydrophobicity since RP-K^SP^ is more hydrophilic than YP-K^SP^ and thus prefers to be exposed to the coacervates-water interface. The magnitude of the peaks in the proximal radial distribution function (pRDF) profiles for water molecules relative to the peptide surface supports these findings (Figure S6b). Consequently, RP-K^SP^/YP-K^SP^ coacervates occupy a slightly larger volume and possess a greater surface area compared to YP-K^SP^ coacervates, suggesting enhanced hydration (Figures S6c and S6d).

The different microenvironments in the coacervates of RP-K^SP^ and YP-K^SP^ could influence the chemical cleavage of K^SP^ modifications, which is triggered by the reduction of its disulfide bond by GSH. As shown in Figures 3f-g, both RP-K^SP^ and YP-K^SP^ peptides in the complex coacervates (formed at 1:1 ratio) exhibited a concentration decay in the presence of GSH (as measured by HPLC), which was well-fitted by first-order reaction kinetics. However, RP-K^SP^ had a 3.2-fold higher reaction rate coefficient compared to YP-K^SP^. This indicates that the reduction rate of the disulfide bond in the K^SP^ side chain, and consequently the disassembly of the complex coacervates, can be tuned by adjusting the RP-K^SP^ to YP-K^SP^ ratio, thereby enabling controlled kinetics of cargo release concomitant with the disassembly of coacervates.

### Intracellular Delivery and Release of Macromolecules Mediated by Complex Coacervates

Our previous studies have demonstrated that homotypic coacervates hold significant potential for the intracellular delivery of macromolecular therapeutics.^[30-31, 49]^ With our understanding of the HB*pep* variants and their derivatives that can form complex coacervates, we further explored their applications as intracellular delivery vehicles for functional macromolecules. We first tested the delivery of EGFP proteins into HeLa cells. Unlike typical complex coacervates driven by electrostatic interactions, which often have limited protein release capabilities,^[50-51]^ the complex coacervates formed by RP and YP at the ratio of 1:1 and 2:1, without the self-immolative K^SP^ modification, successfully delivered and released EGFP within 4 hours, as evidenced by well-distributed fluorescence signals throughout the cells (Figure 4a). However, decreasing the relative YP content resulted in low cell uptake and decreased mean fluorescence intensity (MFI) (Figure 4b), suggesting that the complex coacervates became unstable and prematurely disassembled before entering the cells. We also tested coacervates formed by the mixture of RP and YP-K^SP^, which exhibited high uptake rate but low MFI (Figure 4b). We attribute this result to the increased hydrophobicity of YP-K^SP^ compared to YP, which stabilizes the coacervates but also hinders efficient disassembly and cargo release.

**Figure 4.**
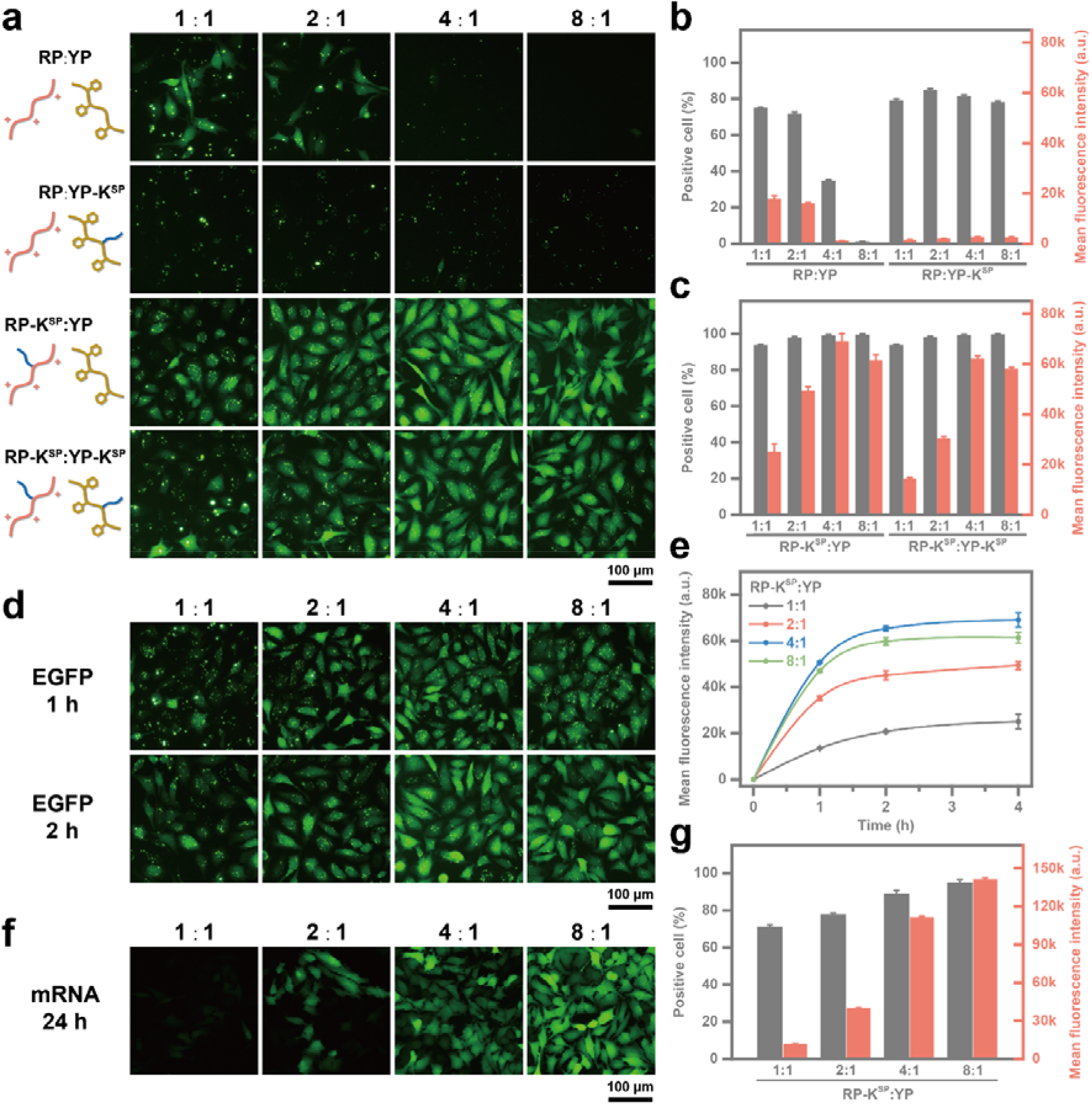
Delivery and release of macromolecules from complex coacervates stabilized by cation-π interactions. **(a-c)** Fluorescence micrographs **(a)** and FACS measurements **(b-c)** of HeLa cells treated with EGFP-loaded complex coacervates for 4 hours. **(d)** Fluorescence micrographs of HeLa cells treated with EGFP-loaded complex coacervates formed by RP-K^SP^/YP with various ratios for 1 and 2 hours. **(e)** Mean fluorescence intensity *vs* time from 0 to 4 hours due to EGFP release from complex coacervates formed by RP-K^SP^/YP at various ratios measured by FACS. Data are presented as the mean □±□ SD of *n* = 3 independent experiments. **(f-g)** Fluorescence micrographs **(f)** and FACS measurements **(g)** of HeLa cells treated with EGFP mRNA-loaded complex coacervates formed by RP-K^SP^/YP at various ratios for 24 hours. Data are presented as the mean □±□ SD of *n* = 3 independent experiments.

We posit that the release of cargo proteins from RP/YP complex coacervates is likely due to the disruption of cation-π interactions by the high concentration of proteins in the crowded cellular environment, which competitively interact with RP and YP (Figure 5a). To test this hypothesis, turbidity measurements were employed to evaluate coacervates formation in the presence of bovine serum albumin (BSA) to mimic the cytosolic environment. As shown in Figures 5b-c, at a low BSA concentration (0.01%), the change in turbidity with peptide concentration was similar to that of the control group without BSA, suggesting that low concentrations of protein cargos do not significantly affect complex coacervation or cargo recruitment. However, when the BSA concentration reached 10%, similar to the total protein concentration inside cells,^[52-53]^ the turbidity of both RP/YP and RP-K^SP^/YP-K^SP^ coacervates significantly decreased, reaching nearly 0% at the peptide concentration of 200 μM. This suggests that the cationic and aromatic peptide variants cannot form coacervates effectively under protein-rich conditions. These findings imply that compared to electrostatic interactions, cation-π interactions may offer a more efficient mechanism to formulate complex coacervates for macromolecule delivery. While the disruption of charge interactions typically requires high salinity,^[54-55]^ which can affect the osmotic pressure of cells and biological systems, cation-π interactions are disrupted by the inherently protein-rich environment of cells.^[56-57]^

**Figure 5.**
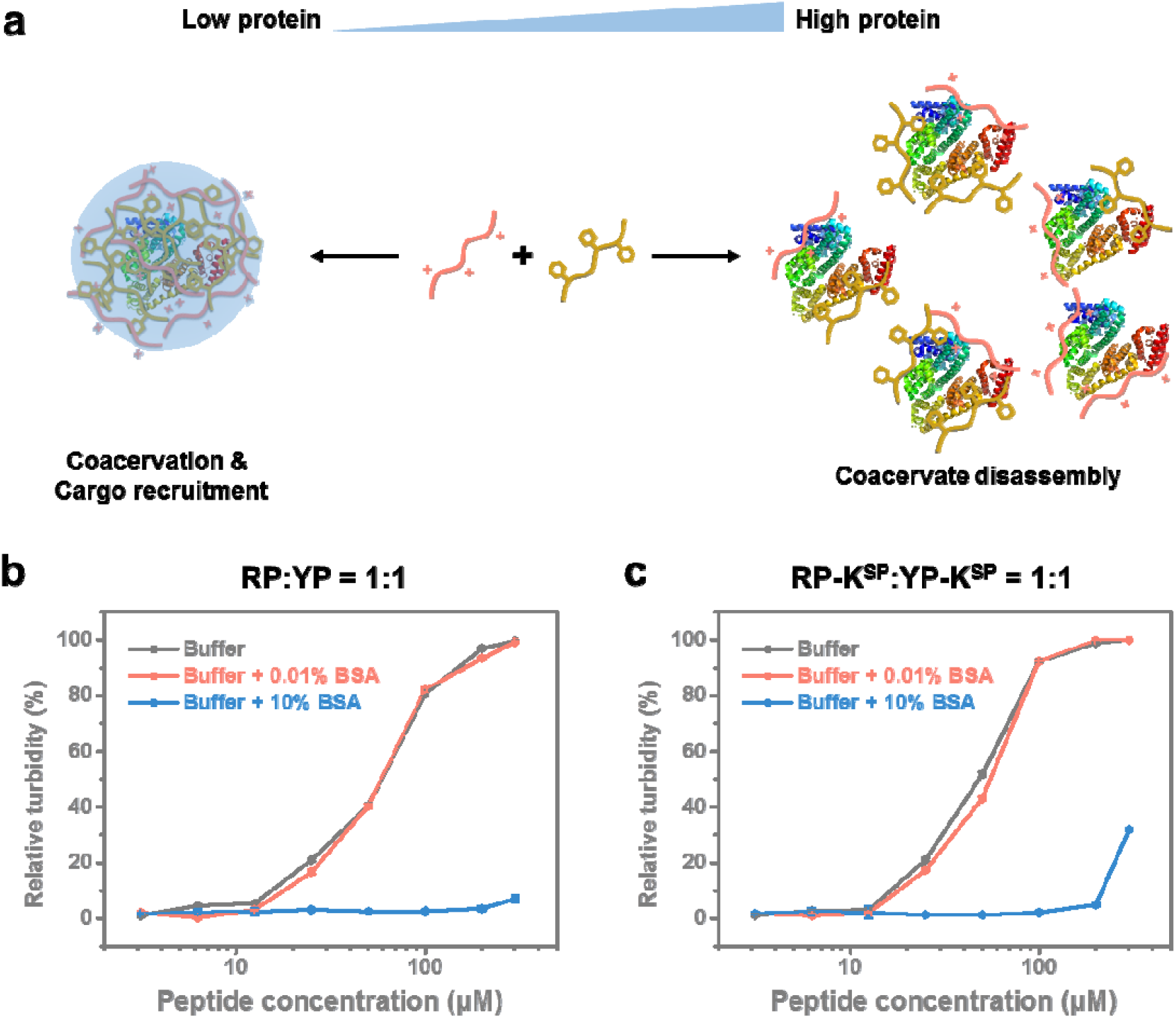
The responsivity of cation-π stabilized complex coacervates to different concentrations of proteins. **(a)** Schematic illustrations of cationic and aromatic peptides interact in low and high concentrations of external proteins. At low protein concentrations, proteins are recruited within the network formed by peptides. However, at high protein concentrations, the protein competes to interact with peptides, disrupting the formation of peptide clusters and further coacervation. **(b-c)** Turbidity measurements of RP:YP = 1:1 **(a)** and RP-K^SP^:YP-K^SP^ = 1:1 **(b)** at zero, low (0.01%) and high (10%) protein concentrations.

Although the RP/YP system demonstrated potential for protein delivery and release, its efficiency was not satisfactory and was limited to a narrow RP/YP ratio range. To address this, the mixtures of RP-K^SP^ with either YP or YP-K^SP^ were tested, as the K^SP^ modification introduces additional hydrophobicity to the cationic peptide, potentially enhancing the stability of the complex coacervates without compromising cargo release. As shown in Figures 4a and c, using RP-K^SP^ as the cationic peptide significantly improved both uptake efficiency and cargo release. Within just 4 hours, HeLa cells treated with EGFP-loaded RP-K^SP^/YP or RP-K^SP^/YP-K^SP^ coacervates exhibited over 90% uptake, along with higher MFI compared to the best-performing RP/YP coacervates. This excellent improvement surpassed our previous efforts using self-coacervating peptides with the same self-immolative modification, which only achieved substantial cargo release after 24 hours.^[31]^ The accelerated release observed here may be attributed to both the chemical cleavage of the self-immolative modification and the responsiveness of cation-π interactions in the crowded cellular environment. Both RP-K^SP^/YP and RP-K^SP^/YP-K^SP^ coacervates showed increased MFI with a rising RP-K^SP^ ratio, peaking at an optimal 4:1 ratio before slightly decreasing due to coacervate instability caused by excessive cationic peptides (Figure 4c). The improvement in delivery efficiency is likely driven by the change of the hydrophobic core formed by aromatic peptides. Similar results were observed when comparing RP-K^SP^/YP and RP-K^SP^/YP-K^SP^ at the same cationic/aromatic peptide ratio, where RP-K^SP^/YP system performed better. Due to its weaker hydrophobicity, YP allows the complex coacervates to have improved access to GSH and proteins in the crowded cellular environment, which serve as triggers for cargo release.

The unique properties of this complex coacervate system provide a potential strategy to control cargo release kinetics simply by adjusting the mixing ratio of the two peptides. To further explore this idea, we conducted a detailed evaluation of the release profiles of complex coacervates formed with various cationic/aromatic peptide ratios, focusing on the best-performing RP-K^SP^/YP system. As shown in Figures 4d-e, cells treated with coacervates at lower RP-K^SP^/YP ratios, such as 1:1 and 2:1, exhibited relatively slower release rates, with EGFP continuously released over the 4-hour measurement period. In contrast, coacervates with higher RP-K^SP^ content reached maximum release within 2 hours, indicating full release of EGFP, which is also evidenced by the disappearance of fluorescence puncta inside the cytoplasm (Figure 4d).

In addition to proteins, the complex coacervate system can also deliver mRNA, another macromolecular therapeutic that has gained significant attention, particularly for its role in the development of COVID-19 vaccines.^[58-59]^ As shown in Figures 4f-g, the RP-K^SP^/YP coacervates with varying mix ratios effectively transfected HeLa cells with a reporter mRNA encoding EGFP, demonstrating a similar pattern to protein delivery: transfection efficiency increased as the YP content decreased. The coacervates achieved the highest transfection efficiency at an 8:1 ratio, successfully transfecting 95% of treated HeLa cells. This robust and versatile delivery capability of the complex coacervates offers significant potential for various applications, including cell therapy, vaccine development, and drug screening.

### Macrophage Engineering Mediated by Complex Coacervates

Macrophage cells are increasingly studied for their pivotal roles in disease-related bioactivities, such as immune response, cancer development, and inflammation.^[60-62]^ However, their intrinsic resistance to foreign material transfections poses a significant challenge in developing targeted therapeutics.^[63]^ To overcome this drawback, we explored the potential of RP-K^SP^/YP coacervates for delivering various functional macromolecules into macrophages. Using EGFP as a model protein, all coacervates with RP-K^SP^/YP ratios ranging from 1:1 to 8:1 successfully transfected over 90% of RAW264.7 macrophages, significantly outperforming the 23.3% transfection efficiency achieved using PULSin, a commercially available reagent commonly used for protein delivery (Figure 6a). Additionally, RP-K^SP^/YP (4:1) coacervates effectively delivered high molecular weight proteins including Alexa Fluor 488-labeled Immunoglobulin G (150 kDa) and R-phycoerythrin (250 kDa), achieving over 99% delivery efficiency (Figures 6b-c). Moreover, the RP-K^SP^/YP coacervates demonstrated excellent mRNA transfection capabilities in RAW264.7 cells, particularly at an 8:1 mix ratio, achieving 83.4% efficiency, significantly higher than the 22.8% efficiency obtained using the highly optimized Lipofectamine MessengerMax reagent (Figure 6d).

**Figure 6.**
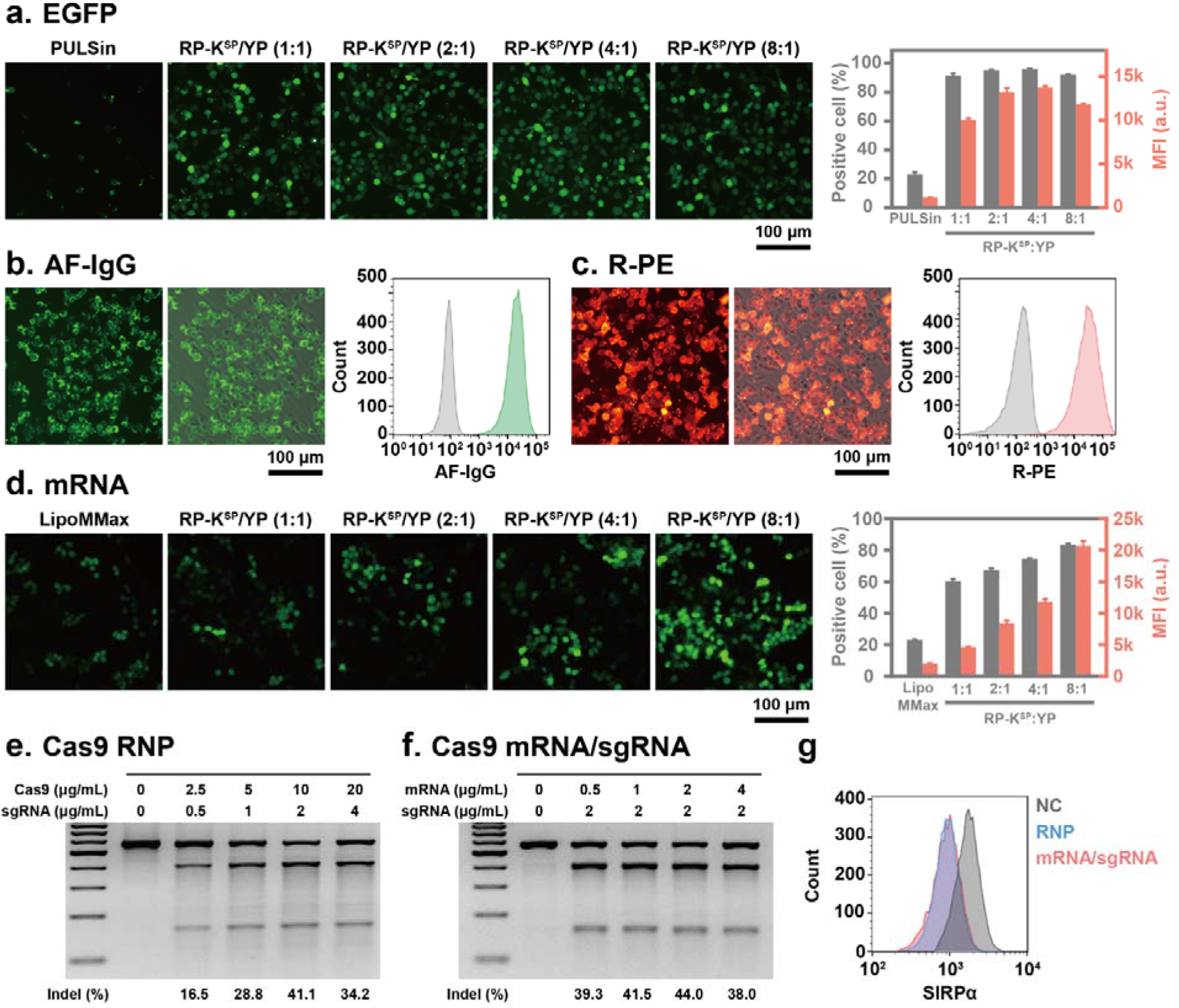
Macrophage engineering mediated by RP-K^SP^/YP complex coacervates. **(a)** Fluorescence micrographs and FACS measurements of RAW264.7 cells treated with EGFP-loaded complex coacervates formed by RP-K^SP^/YP with various ratios for 4 hours compared to the commercial reagent PULSin. Data are presented as the mean □±□ SD of *n* = 3 independent experiments. **(b)** Fluorescence micrographs (left), merged images of the fluorescence channel and optical widefield channel (middle), and FACS analysis (right) of RAW264.7 cells treated with AF-IgG-loaded complex coacervates formed by RP-K^SP^:YP = 4:1 for 4 hours. **(c)** Fluorescence micrographs (left), merged images of the fluorescence channel and optical widefield channel (middle), and FACS analysis (right) of RAW264.7 cells treated with R-PE-loaded complex coacervates formed by RP-K^SP^:YP = 4:1 for 4 hours. **(d)** Fluorescence micrographs and FACS measurements of RAW264.7 cells treated with EGFP mRNA-loaded complex coacervates formed by RP-K^SP^/YP with various ratios for 24 hours compared to the commercial reagent Lipofectamine MessengerMax (MMax). Data are presented as the mean □±□ SD of *n* = 3 independent experiments. **(e-f)** Analysis of indel frequency at the SIRPα locus in RAW264.7 cells treated with SIRPα-targeted Cas9 RNP-loaded complex coacervates formed by RP-K^SP^:YP = 4:1 (e) and SIRPα-targeted Cas9 mRNA/sgRNA-loaded complex coacervates formed by RP-K^SP^:YP = 8:1 (f). **(g)** The expression of SIRPα on edited RAW264.7 cells measured by FACS compared to the negative control (NC) group, which did not undergo editing.

Encouraged by the promising performance of our complex coacervates delivery system, we then explored the delivery of more complex therapeutics, namely the clustered regularly interspaced short palindromic repeats (CRISPR)/CRISPR associated protein 9 (Cas9) genome editing tools. The delivery of CRISPR/Cas9 tools is considered challenging as it involves delivering two components: the Cas9 nuclease and single guided RNA (sgRNA).^[64]^ The most common way to tackle this is to deliver an all-in-one plasmid that encodes both the sequence of Cas9 and sgRNA.^[65]^ However, plasmid based editing tools are associated with risks of genome integration and off-target effects.^[66]^ To mitigate these risks, we employed RP-K^SP^/YP coacervates for the delivery of two plasmid-free CRISPR/Cas9 tools: a Cas9 mRNA/sgRNA mixture and Cas9/sgRNA ribonucleoprotein (RNP), both targeting the signal-regulatory protein alpha (SIRPα) gene. SIRPα is a cell surface receptor primarily expressed on macrophages that interacts with CD47 to deliver a “don’t eat me” signal, preventing macrophages from engulfing healthy cells. However, many tumor cells overexpress CD47 to evade immune detection, making the SIRPα-CD47 axis a prime target for cancer immunotherapies. Disrupting this interaction can enhance macrophage-mediated elimination of cancer cells.^[33-34]^ The T7 Endonuclease I (T7EI) assay confirmed successful editing at the SIRPα locus, with insertion-deletion (indel) frequencies exceeding 40% at optimal cargo concentrations (Figures 6e-f). Additionally, flow cytometry analysis demonstrated a significant decrease in SIRPα expression in the edited cells (Figure 6g), highlighting the potential of coacervate-mediated delivery systems for macrophage cell therapies.

## Conclusion

This study presents a novel strategy for designing a complex coacervate-based intracellular delivery system using the HB*pep* family of peptides. By incorporating cationic, anionic, and aromatic residues into the HB*pep* sequence, we systematically investigated how these substitutions affect both self- and complex coacervation behaviors. Our findings indicate that a low density of oppositely charged residues is unable to establish stable electrostatic interactions, resulting in homogeneous solutions of KP and EP and failure to form complex coacervates. Conversely, the strong cation-π interactions between RP and YP prevent RP dissolution and YP aggregation, instead promoting and stabilizing the formation of complex coacervates. Notably, the cation-π interactions are disrupted by the protein-rich cellular environment, serving as a trigger for coacervate disassembly and cargo release in cells.

By introducing Lys and a self-immolative disulfide moiety into the peptide sequences, we engineered complex coacervates capable of responding to intracellular redox conditions, thereby enhancing the controlled release of cargos. Adjusting the ratio of cationic to aromatic peptides allowed precise control over the phase behavior and release kinetics of cargos from the coacervates, facilitating the efficient delivery of a diverse range of macromolecular therapeutics, including proteins, mRNA, and CRISPR/Cas9 gene editing tools. Notably, this approach proved effective at genetically editing macrophages, a cell type notoriously resistant to transfection, highlighting the versatility and adaptability of our peptide-based complex coacervate system for immune cell therapies.

Our findings emphasize the importance of cation-π interactions in designing complex coacervates with tunable stability and cargo release profiles, offering a new avenue for the development of advanced intracellular drug delivery vehicles. This strategy not only enhances the performance of coacervates as delivery vehicles but also offers potential solutions to broader biomedical challenges, such as those related to protein aggregation. This work lays the foundation for future explorations of peptide-based coacervates in chemical biology, therapeutic delivery, and beyond, with significant implications for drug development, vaccine design, and cell therapy applications.

## Supporting Information

Experimental section and supplementary data are described in detail in the Supporting Information. The authors have cited additional references within the Supporting Information.

## Acknowledgments

This research was funded by the Singapore Ministry of Education (MOE) through an Academic Research Fund (AcRF) Tier 3 grant (grant no. MOE 2019-T3-1-012) and the Singapore National Research Foundation (NRF) though its Competitive Research Program (grant no. NRF-CRP30-2023-0004). We thank the Bioinformatics Institute (A*STAR) and the National Supercomputing Centre Singapore for computational support.

## References

[1] A. Miserez, J. Yu, P. Mohammadi, Chem. Rev. 2023, 123, 2049–2111.

[2] M. Corley, M. C. Burns, G. W. Yeo, Mol. Cell 2020, 78, 9–29.

[3] B. K. Ahn, J. Am. Chem. Soc. 2017, 139, 10166–10171.

[4] D. T. Moore, B. W. Berger, W. F. DeGrado, Structure 2008, 16, 991–1001.

[5] G. Bao, S. Suresh, Nat. Mater. 2003, 2, 715–725.

[6] M. J. Harrington, R. Mezzenga, A. Miserez, Nature Reviews Bioengineering 2024, 2, 260–278.

[7] A. Sathyavageeswaran, J. Bonesso Sabadini, S. L. Perry, Acc. Chem. Res. 2024, 57, 386–398.

[8] M. Abbas, W. P. Lipiński, J. Wang, E. Spruijt, Chem. Soc. Rev. 2021, 50, 3690–3705.

[9] J. Dinic, A. B. Marciel, M. V. Tirrell, Curr. Opin. Colloid Interface Sci. 2021, 54, 101457.

[10] N. Doshi, W. Guo, F. Chen, P. Venema, H. C. Shum, R. de Vries, X. Li, Soft Matter 2024, 20, 1966–1977.

[11] Q. Peng, T. Wang, D. Yang, X. Peng, H. Zhang, H. Zeng, Prog. Polym. Sci. 2024, 153, 101827.

[12] S. P. O. Danielsen, J. McCarty, J.-E. Shea, K. T. Delaney, G. H. Fredrickson, Proceedings of the National Academy of Sciences 2019, 116, 8224–8232.

[13] P. Mohammadi, G. Beaune, B. T. Stokke, J. V. I. Timonen, M. B. Linder, ACS Macro Letters 2018, 7, 1120–1125.

[14] F. G. Quiroz, A. Chilkoti, Nat. Mater. 2015, 14, 1164–1171.

[15] D. López Barreiro, I. J. Minten, J. C. Thies, C. M. J. Sagt, ACS Biomaterials Science & Engineering 2023, 9, 3796–3809.

[16] A. K. Varanko, J. C. Su, A. Chilkoti, Annu. Rev. Biomed. Eng. 2020, 22, 343–369.

[17] C. E. Sing, S. L. Perry, Soft Matter 2020, 16, 2885–2914.

[18] Y. P. Timilsena, T. O. Akanbi, N. Khalid, B. Adhikari, C. J. Barrow, Int. J. Biol. Macromol. 2019, 121, 1276–1286.

[19] C. G. de Kruif, F. Weinbreck, R. de Vries, Curr. Opin. Colloid Interface Sci. 2004, 9, 340–349.

[20] J. K. A. Tom, A. A. Deniz, Curr. Opin. Colloid Interface Sci. 2021, 56, 101488.

[21] C. Zhang, H. Peng, J. H. Waite, Q. Zhao, J. Am. Chem. Soc. 2024, 146, 2219–2226.

[22] Y. Sun, S. H. Hiew, A. Miserez, Acc. Chem. Res. 2024, 57, 164–174.

[23] B. Gabryelczyk, H. Cai, X. Shi, Y. Sun, P. J. M. Swinkels, S. Salentinig, K. Pervushin, A. Miserez, Nat. Commun. 2019, 10, 5465.

[24] H. Cai, B. Gabryelczyk, M. S. S. Manimekalai, G. Grüber, S. Salentinig, A. Miserez, Soft Matter 2017, 13, 7740–7752.

[25] Y. Tan, S. Hoon, P. A. Guerette, W. Wei, A. Ghadban, C. Hao, A. Miserez, J. H. Waite, Nat. Chem. Biol. 2015, 11, 488–495.

[26] J. Lim, A. Kumar, K. Low, C. S. Verma, Y. Mu, A. Miserez, K. Pervushin, The Journal of Physical Chemistry B 2021, 125, 6776–6790.

[27] X. Wu, Y. Sun, J. Yu, A. Miserez, Communications Chemistry 2024, 7, 5.

[28] Z. W. Lim, Y. Ping, A. Miserez, Bioconjug. Chem. 2018, 29, 2176–2180.

[29] Z. W. Lim, V. B. Varma, R. V. Ramanujan, A. Miserez, Acta Biomater. 2020, 110, 221–230.

[30] Y. Sun, X. Xu, L. Chen, W. L. Chew, Y. Ping, A. Miserez, ACS Nano 2023, 17, 16597–16606.

[31] Y. Sun, S. Y. Lau, Z. W. Lim, S. C. Chang, F. Ghadessy, A. Partridge, A. Miserez, Nat. Chem. 2022, 14, 274–283.

[32] G. C. Yeo, F. W. Keeley, A. S. Weiss, Adv. Colloid Interface Sci. 2011, 167, 94–103.

[33] M. P. Chao, I. L. Weissman, R. Majeti, Curr. Opin. Immunol. 2012, 24, 225–232.

[34] M. E. W. Logtenberg, F. A. Scheeren, T. N. Schumacher, Immunity 2020, 52, 742–752.

[35] A. M. Rumyantsev, N. E. Jackson, J. J. de Pablo, Annual Review of Condensed Matter Physics 2021, 12, 155–176.

[36] L. Li, S. Srivastava, M. Andreev, A. B. Marciel, J. J. de Pablo, M. V. Tirrell, Macromolecules 2018, 51, 2988–2995.

[37] S. L. Perry, L. Leon, K. Q. Hoffmann, M. J. Kade, D. Priftis, K. A. Black, D. Wong, R. A. Klein, C. F. Pierce, K. O. Margossian, J. K. Whitmer, J. Qin, J. J. de Pablo, M. Tirrell, Nat. Commun. 2015, 6, 6052.

[38] Y. Hong, S. Najafi, T. Casey, J.-E. Shea, S.-I. Han, D. S. Hwang, Nat. Commun. 2022, 13, 7326.

[39] A. A. Polyansky, L. D. Gallego, R. G. Efremov, A. Köhler, B. Zagrovic, eLife 2023, 12, e80038.

[40] J. P. Gallivan, D. A. Dougherty, Proceedings of the National Academy of Sciences 1999, 96, 9459–9464.

[41] J. N. Israelachvili, K. Kristiansen, M. A. Gebbie, D. W. Lee, S. H. Donaldson, Jr., S. Das, M. V. Rapp, X. Banquy, M. Valtiner, J. Yu, The Journal of Physical Chemistry B 2013, 117, 16369–16387.

[42] J. Yu, N. E. Jackson, X. Xu, Y. Morgenstern, Y. Kaufman, M. Ruths, J. J. de Pablo, M. Tirrell, Science 2018, 360, 1434–1438.

[43] D. S. Hwang, H. Zeng, A. Srivastava, D. V. Krogstad, M. Tirrell, J. N. Israelachvili, J. H. Waite, Soft Matter 2010, 6, 3232–3236.

[44] Q. Guo, G. Zou, X. Qian, S. Chen, H. Gao, J. Yu, Nat. Commun. 2022, 13, 5771.

[45] X. Han, G. Li, G. Li, K. Lin, Biochemistry 1998, 37, 10730–10737.

[46] Y. Lin, W. Li, J. Wu, H. Zhang, R. W. Colman, Thromb. Res. 1998, 90, 65–72.

[47] M. R. Ajmal, in Diseases, Vol. 11, 2023.

[48] N. Gregersen, L. Bolund, P. Bross, Mol. Biotechnol. 2005, 31, 141–150.

[49] J. Liu, E. Spruijt, A. Miserez, R. Langer, Nature Reviews Materials 2023, 8, 139–141.

[50] W. Xiao, M. D. Jakimowicz, I. Zampetakis, S. Neely, F. Scarpa, S. A. Davis, D. S. Williams, A. W. Perriman, Advanced Biosystems 2020, 4, 2000101.

[51] K. A. Black, D. Priftis, S. L. Perry, J. Yip, W. Y. Byun, M. Tirrell, ACS Macro Letters 2014, 3, 1088–1091.

[52] A. T. Molines, J. Lemière, M. Gazzola, I. E. Steinmark, C. H. Edrington, C.-T. Hsu, P. Real-Calderon, K. Suhling, G. Goshima, L. J. Holt, M. Thery, G. J. Brouhard, F. Chang, Dev. Cell 2022, 57, 466–479.e466.

[53] T. Kühn, T. O. Ihalainen, J. Hyväluoma, N. Dross, S. F. Willman, J. Langowski, M. Vihinen-Ranta, J. Timonen, PLoS ONE 2011, 6, e22962.

[54] S. L. Perry, Y. Li, D. Priftis, L. Leon, M. Tirrell, in Polymers, Vol. 6, 2014, pp. 1756–1772.

[55] W. C. Blocher, S. L. Perry, WIREs Nanomedicine and Nanobiotechnology 2017, 9, e1442.

[56] D. A. Dougherty, The Journal of Nutrition 2007, 137, 1504S–1508S.

[57] A. R. Subramanya, C. R. Boyd-Shiwarski, Annu. Rev. Physiol. 2024, 86, 429–452.

[58] L. R. Baden, H. M. El Sahly, B. Essink, K. Kotloff, S. Frey, R. Novak, D. Diemert, S. A. Spector, N. Rouphael, C. B. Creech, J. McGettigan, S. Khetan, N. Segall, J. Solis, A. Brosz, C. Fierro, H. Schwartz, K. Neuzil, L. Corey, P. Gilbert, H. Janes, D. Follmann, M. Marovich, J. Mascola, L. Polakowski, J. Ledgerwood, B. S. Graham, H. Bennett, R. Pajon, C. Knightly, B. Leav, W. Deng, H. Zhou, S. Han, M. Ivarsson, J. Miller, T. Zaks, N. Engl. J. Med. 2020.

[59] F. P. Polack, S. J. Thomas, N. Kitchin, J. Absalon, A. Gurtman, S. Lockhart, J. L. Perez, G. Pérez Marc, E. D. Moreira, C. Zerbini, R. Bailey, K. A. Swanson, S. Roychoudhury, K. Koury, P. Li, W. V. Kalina, D. Cooper, R. W. Frenck, L. L. Hammitt, Ö. Türeci, H. Nell, A. Schaefer, S. Ünal, D. B. Tresnan, S. Mather, P. R. Dormitzer, U. Şahin, K. U. Jansen, W. C. Gruber, N. Engl. J. Med. 2020, 383, 2603–2615.

[60] A. Chawla, K. D. Nguyen, Y. P. S. Goh, Nat. Rev. Immunol. 2011, 11, 738–749.

[61] S. Chen, A. F. U. H. Saeed, Q. Liu, Q. Jiang, H. Xu, G. G. Xiao, L. Rao, Y. Duo, Signal Transduction and Targeted Therapy 2023, 8, 207.

[62] A. Mantovani, P. Allavena, F. Marchesi, C. Garlanda, Nat. Rev. Drug Discovery 2022, 21, 799–820.

[63] M. Klichinsky, M. Ruella, O. Shestova, X. M. Lu, A. Best, M. Zeeman, M. Schmierer, K. Gabrusiewicz, N. R. Anderson, N. E. Petty, K. D. Cummins, F. Shen, X. Shan, K. Veliz, K. Blouch, Y. Yashiro-Ohtani, S. S. Kenderian, M. Y. Kim, R. S. O’Connor, S. R. Wallace, M. S. Kozlowski, D. M. Marchione, M. Shestov, B. A. Garcia, C. H. June, S. Gill, Nat. Biotechnol. 2020, 38, 947–953.

[64] C. A. Lino, J. C. Harper, J. P. Carney, J. A. Timlin, Drug Delivery 2018, 25, 1234–1257.

[65] H.-X. Wang, M. Li, C. M. Lee, S. Chakraborty, H.-W. Kim, G. Bao, K. W. Leong, Chem. Rev. 2017, 117, 9874–9906.

[66] Z. Liang, K. Chen, T. Li, Y. Zhang, Y. Wang, Q. Zhao, J. Liu, H. Zhang, C. Liu, Y. Ran, C. Gao, Nat. Commun. 2017, 8, 14261.

