## Supplementary information for "Peptide-Based Complex Coacervates Stabilized by Cation-π Interactions for Cell Engineering"

**Experimental Section**

**Materials**

Resins and fluorenylmethoxycarbonyl (Fmoc)-protected amino acids used in solid-phase peptide synthesis were purchased from GL Biochem. N-hydroxysuccinimide, tetrahydrofuran, N,N′-diisopropylcarbodiimide, triphosgene and benzoic acid were purchased from Tokyo Chemical Industry (TCI). N,N-diisopropylethylamine, 2-hydroxyethyl disulfide, piperidine, trifluoroacetic acid, triisopropylsilane, R-phycoerythrin, Oxyma, and reduced L-glutathione (GSH) were obtained from Sigma-Aldrich. Dichloromethane, N,N-dimethylformamide (DMF), Alexa Fluor 488 NHS ester, SYBR Safe DNA gel stain, Opti-MEM, Alexa Fluor 488 labelled anti-mouse SIRPα antibody, and Lipofectamine MessengerMAX Reagent were purchased from Thermo Fisher Scientific. Cell counting kit-8 (CCK-8) was purchased from Abcam. PULSin protein transfection reagent was purchased from Polyplus. T7 Endonuclease I, Q5 Hot Start high-fidelity 2X master mix, and 100 bp DNA Ladder were purchased from New England Biolabs. Organic solvents, including ethyl acetate, hexane and diethyl ether were purchased from Aik Moh Paints & Chemicals Pte Ltd. Dulbecco’s modified Eagle medium (DMEM), fetal bovine serum (FBS), phosphate-buffered saline (PBS) and antibiotic-antimycotic (100X) liquid were purchased from Gibco. EGFP-encoding mRNA, Cas9-encoding mRNA, SIRPα targeting sgRNA and primers for amplifying SIRP locus were obtained from GenScript. HeLa and RAW264.7 cell lines were obtained from ATCC.

**Peptide synthesis and modification**

The HB*pep* variants were synthesized on a microwave-assisted solid phase peptide synthesizer (Liberty Blue) using N,N′-diisopropylcarbodiimide (DIC)/Oxyma as coupling reagents and 20% piperidine in DMF as deprotection reagents. After the synthesis, peptides were cleaved from the resins using a cocktail containing 95% of trifluoroacetic acid (TFA), 2.5% of H_2_O and 2.5% of triisopropylsilane (TIPS) for 2 h. Then the supernatants were collected by filtration and concentrated using a nitrogen flow, followed by precipitating into 50 mL of cold diethyl ether. The pellets from centrifugation were dried under vacuum and re-dissolved using 5% acetic acid aqueous solution for purification by HPLC (1260 Infinity, Agilent Technologies) equipped with a C8 column (Zorbax 300SB-C8, Agilent Technologies). The purified Fmoc-HB*pep* variants were isolated by lyophilization from HPLC elutes.

The HB*pep*-K^SP^ variants prepared by modifying the Lys side chain of the N-terminus protected Fmoc-HB*pep*-K peptide, followed by the Fmoc deprotection. First, the Fmoc-HB*pep*-K backbone was prepared by peptide synthesizer as mentioned above. The Lys side chain amine of Fmoc-HB*pep*-K then reacts with an amine-reactive small molecule NHS-SS-Ph (Figure S1), followed by deprotection to produce HB*pep*-K^SP^ as described in our previous works.^[1-2]^ In detail, 15 μmol of peptides was dissolved in 3 mL of dimethylformamide (DMF). Then, 450 μmol of N,N-diisopropylethylamine (DIEA) was added into the peptide solution followed by 100 μL of DMF containing 20 μmol of NHS-SS-Ph. After overnight reaction at room temperature, 1 mL of piperidine was added into the mixture for another 1 h of Fmoc deprotection. The raw products were precipitated out by adding 30 mL of cold diethyl ether and collected by centrifugation. The pellets were dried under vacuum and re-dissolved using 5% acetic acid aqueous solution for purification by HPLC equipped with a C8 reverse phase column. The purified HB*pep*-K^SP^ variants were isolated by lyophilization from the HPLC elutes, whose MW was confirmed by Matrix Assisted Laser Desorption/Ionization (MALDI) time-of-flight (ToF) mass spectrometry (AXIMA Performance spectrometer, Shimadzu) using α-cyano-4-hydroxycinnamic acid (CHCA) as the matrix, shown in Figure S7.

**LLPS of peptide variants and their mixture**

The phase diagram of variants and mixtures was determined by observing the phase separation at different pHs and concentrations via an inverted microscope (AxioObserver.Z1, Zeiss). Peptides were dissolved in 10 mM acetic acid aqueous solution with various concentrations as stock solutions. The LLPS was induced by mixing the stock solutions with buffers at a volume ratio of 1:9. The buffer details are described in Table S1.

To further confirm the presence of cation-π interactions between peptides, YP-K^SP^ aggregates were pre-formed by mixing 10 μL of stock solution (3 mM) with 90 μL of buffer (pH 7.0, IS = 100 mM). The mixture was transferred onto a glass slide and incubated for 5 minutes to allow the aggregates to settle. Afterward, the supernatant was replaced with 90 μL of fresh buffer (pH 7.0, IS = 100 mM), followed by the addition of 10 μL of RP-K^SP^ stock solution (3 mM). The mixture was then incubated at room temperature and imaged at various time points using an inverted microscope (AxioObserver.Z1, Zeiss).

**Fourier-transform infrared (FTIR) spectroscopy**

FTIR spectroscopy was employed to identify the secondary structure of peptide coacervates and aggregates. The spectra were collected at room temperature in the attenuated total reflection (ATR) mode on a FTIR spectrometer (Vertex 70, Bruker). Before the signal acquisition, 100 μL of suspensions were prepared by mixing 10 μL of stock solution (3 mM in D_2_O containing 10 mM acetic acid) with 90 μL of buffers (10 mM phosphate D_2_O buffer with 7.0 of pH and 100 mM of ionic strength fixed by NaCl), and then transferred to the surface of the ZnSe-diamond in the ATR accessory. The amide I band range from 1800 to 1350 cm^-1^ was collected and then processed to substrate water vapors and black sample, and correct baselines by OPUS 6.5 software.^[3-4]^

**Surface force apparatus (SFA) measurements**

Both the viscoelastic properties of complex coacervates and the interactions between two peptide layers was investigated using an SFA 2000 (SurForce LLC, Santa Barbara).^[5]^ As described in previous studies,^[6-7]^ freshly cleaved mica with a 55 nm silver layer deposited on its back was glued on the glass disks. For coacervate samples, 2 μL of freshly prepared CMs (0.3 mM in buffer, pH 7.0) was mixed with 18 μL of PBS and injected in the gap between two mica surfaces. The distance *D* between two surfaces was measured and calculated based on the fringes of equal chromatic order (FECO) technique. After the sample was injected into the gap, the system was equilibrated for 30 min by keeping two surfaces in contact with a bridging coacervate film. Then, the two surfaces started to approach followed by separation. The measured force *F* was normalized by the effective radius of the surface *R*.

To measure the interactions between two peptide layers, 100 μL of peptide solution (15 μM in 10 mM acetic acid) was applied to a mica-coated glass disk and incubated for 20 minutes at room temperature. The mica was then rinsed with the pH 7.0 buffer and assembled into the SFA 2000. Between the two mica surfaces, 20 μL of pH 7.0 buffer, with or without 100 mM TMA, was introduced. The system was equilibrated for 30 minutes, allowing the two surfaces to form a liquid bridge before testing, as described earlier.

**Molecular dynamics (MD) simulations**

To understand how intermolecular interactions modulate the liquid-liquid phase separation behaviors in the peptides, MD simulations were carried out for three systems: pure RP-K^SP^ (60 RP-K^SP^ molecules); a 1:1 mixture of RP-K^SP^ and YP-K^SP^ (30 RP-K^SP^ and 30 YP-K^SP^ molecules) and pure YP-K^SP^ (60YP-K^SP^ molecules). Each system was subject to two replicates of simulations. The peptides were initially randomly placed in the box, followed by solvation with water molecules and neutralization with 0.15 M NaCl. Each system was initially subject to 500 steps using steep descent energy minimization, followed by a 100 ps of MD simulation in the NVT ensemble. To accelerate sampling, the production run is divided into two steps: 500 ns simulations at 500 K followed by another 500 ns of production run at 300 K. The details of each system were summarized in Table S2. The proximal radial distribution functions (pRDF), solvent accessible surface area (SASA), the number of pi-pi and cation-pi interaction pairs were calculated using the combination of the last 300 ns of the two replicates.^[8-9]^ To understand the distributions of RP-K^SP^ and YP-K^SP^ in the cluster, we also performed MD simulations using a slab box.^[10-11]^ The clusters from the above simulations were placed in a slab with the ratio of box length x:y:z=1:1:4. Similar to the simulations using a cubic box, a two-step simulation was carried out for the production runs: a 500 ns simulation at 500 K followed by a 500 ns simulation at at 300K (Table S2). Two replicates of slab simulations were carried out for each system, and the last 300 ns of each simulation was used for analysis.

In all simulations, the peptides were modelled using the AMBER14sb force field with water described by the TIP3P model.^[12-13]^ Parameters of the unnatural amino acid KSP were obtained using the antechamber module of the AMBER 20 package.^[14]^ Lennard-Jones and short-range electrostatic interactions were computed using a cutoff of 0.9 nm, while long-range electrostatic interactions were calculated using PME.^[15]^ All simulations were carried out in the NPT ensemble with temperature and pressure maintained at 300 K and 1 bar. All simulations were carried out using GROMACS 2021 patched with Plumed-2.9.^[16-17]^

**Redox-responsivity evaluations**

The difference in the reduction rate of self-immolative side chain of RP-K^SP^ and YP-K^SP^ in the RP-K^SP^:YP-K^SP^ = 1:1 complex coacervates was evaluated by measuring the concentration decrease of unreacted peptides in the presence of 1 mM GSH. The freshly prepared RP-K^SP^:YP-K^SP^ = 1:1 complex coacervates (100 μL, 0.3 mM) were diluted in 900 μL of PBS containing 1.11 mM of GSH. The mixtures were incubated at 37 °C for different time periods before adding 50 μL of acetic acid to dissolve all the unreacted peptides, and their concentrations were measured by HPLC.

**Cell cultures**

HeLa and RAW 264.7 cells were cultured in DMEM supplemented with 10% FBS, 100 U/mL penicillin, and 100 μg/mL streptomycin under typical conditions (37 °C and 5% CO_2_). Jurkat cells were cultured in RPMI-1640 Medium supplemented with 10% FBS, 100 U/mL penicillin, and 100 μg/mL streptomycin. HeLa-EGFP cells were cultured in DMEM supplemented with 10% FBS, 100 U/mL penicillin, 100 μg/mL streptomycin, and 10 μg/mL blasticidin. For HeLa cells, the subculture started by detaching the cells with trypsin treatment, followed by centrifugation (1000 rpm, 5 min) to collect the cells. Then the pellets were resuspended with fresh media for subculture or experiments. For RAW 264.7 cells, the cells were detached from the culture flask using a cell scraper (Corning) and collected by centrifugation (1000 rpm, 5 min). Then the pellets were resuspended with fresh media for subculture or experiments.

**Protein delivery**

To perform protein deliveries, cells were suspended in 2 mL of full media and transferred into 35 cm^2^ culture dishes. When the confluency reached ~ 60%, the medium was replaced with 900 μL of Opti-MEM, then 100 μL of freshly prepared coacervates (0.3 mM peptides, 0.1 mg/mL proteins) were prepared by adding the peptide stocks into cargos containing buffer (pH 7.0 buffer for RP-K^SP^/YP-K^SP^ and RP-K^SP^/YP, and pH 7.0 buffer for RP/YP and RP/YP-K^SP^), and added into the Opti-MEM. The treated cells were imaged at different time points including 1 h, 2 h, and 4 h using fluorescence microscopy (AxioObserver.Z1, Zeiss), the cellular uptake and cargo release were also quantified by FACS (LSR Fortessa X20, BD Biosciences). The commercially available protein transfection reagent PULSin (Polyplus) was used as a comparison according to protocols from the manufacturers.

**Gene transfection**

To evaluate the gene transfection efficiency of complex coacervates, the mRNA encoding EGFP reporter gene were used as cargos. Before transfection, cells were incubated in 35 cm^2^ dishes until the confluency reached ~ 60%. The medium was then replaced with 900 μL of Opti-MEM, followed by the addition of 100 μL of freshly prepared mRNA-loaded coacervates (0.3 mM peptides, 10 μg/mL mRNA). After 4  h of incubation, the medium was removed, and the cells were washed with PBS twice before adding 2 mL of the full medium. Transfection was then continued for another 20 h before imaging the cells under a fluorescence microscope and testing the transfection efficiency by FACS.

To deliver all three types of CRISPR/Cas9 genome editing modalities, RAW264.7 cells are cultured in 35 cm^2^ dishes until reaching 40% confluence. Then the medium was replaced with 900 μL of Opti-MEM and 100 μL of cargo-loaded coacervates. After 4 h of uptake, the medium was discarded. The cells were washed with PBS twice and cultured in full media for another 44 h. The efficiency of 48 h of transfection can be evaluated by using the T7 Endonuclease 1 (T7EI) assay. First, the genomic DNA was extracted by using a DNeasy blood and tissue kit (QIAGEN). The target genomic locus was amplified by PCR using Q5 Hot Start high-fidelity 2X master mix (NEB) and primers listed in Table S3 and purified by PureLink PCR purification kit (Thermo Fisher Scientific). Then, 200 ng of PCR products were digested by T7EI and analyzed by 2% agarose gels before imaging with the gel documentation system. The gray level of digested bands and undigested bands was measured by ImageJ. The indel percentage could be calculated by the following formula:^[18-19]^

[1−(1−fraction cleaved)^1/2^]×100

where fraction cleaved = the sum of each digested band intensity/(the sum of each digested band intensity + undigested band intensity).

The knock-out efficiency was also quantified using Alexa Fluor 488 labelled anti-mouse SIRPα antibody and FACS.

**Cytotoxicity study**

The cytotoxicity of complex coacervates was evaluated using the Cell Counting Kit-8 (CCK-8). As described previously,^[20]^ cells were cultured in 96-well plates with 100 μL of full medium and incubated for 24  h. The medium was then replaced with 90 μL of Opti-MEM mixed with 10 μL of complex coacervates (0.3 mM peptides). After 4 h of uptake, the medium was removed, and the cells were washed with PBS twice and cultured in 100 μL of fresh full medium. The cells were incubated for another 20 h before changing the medium to the full medium containing 10% CCK-8 solution. After 4 h of incubation, the cells were measured for absorbance at 460 nm using a microplate reader (Infinite M200 Pro, Tecan). The relative cell viability was calculated as:

$$\frac{A_{t}-A_{b}}{A_{c}-A_{b}}\times100\%$$

where *A*_t_, *A*_b_, and *A*_c_ represent the absorbance of tested cells, no cells, and untreated cells, respectively.

All complex coacervates showed negligible cytotoxicity in both HeLa and RAW264.7 cell lines (Figure S8).

**Statistics and reproducibility**

All experiments were repeated three times. The data are presented as mean ± standard deviation (SD). All microscopy experiments were repeated independently three times and the presented images are representative of the obtained data.

**Supplementary Figures**

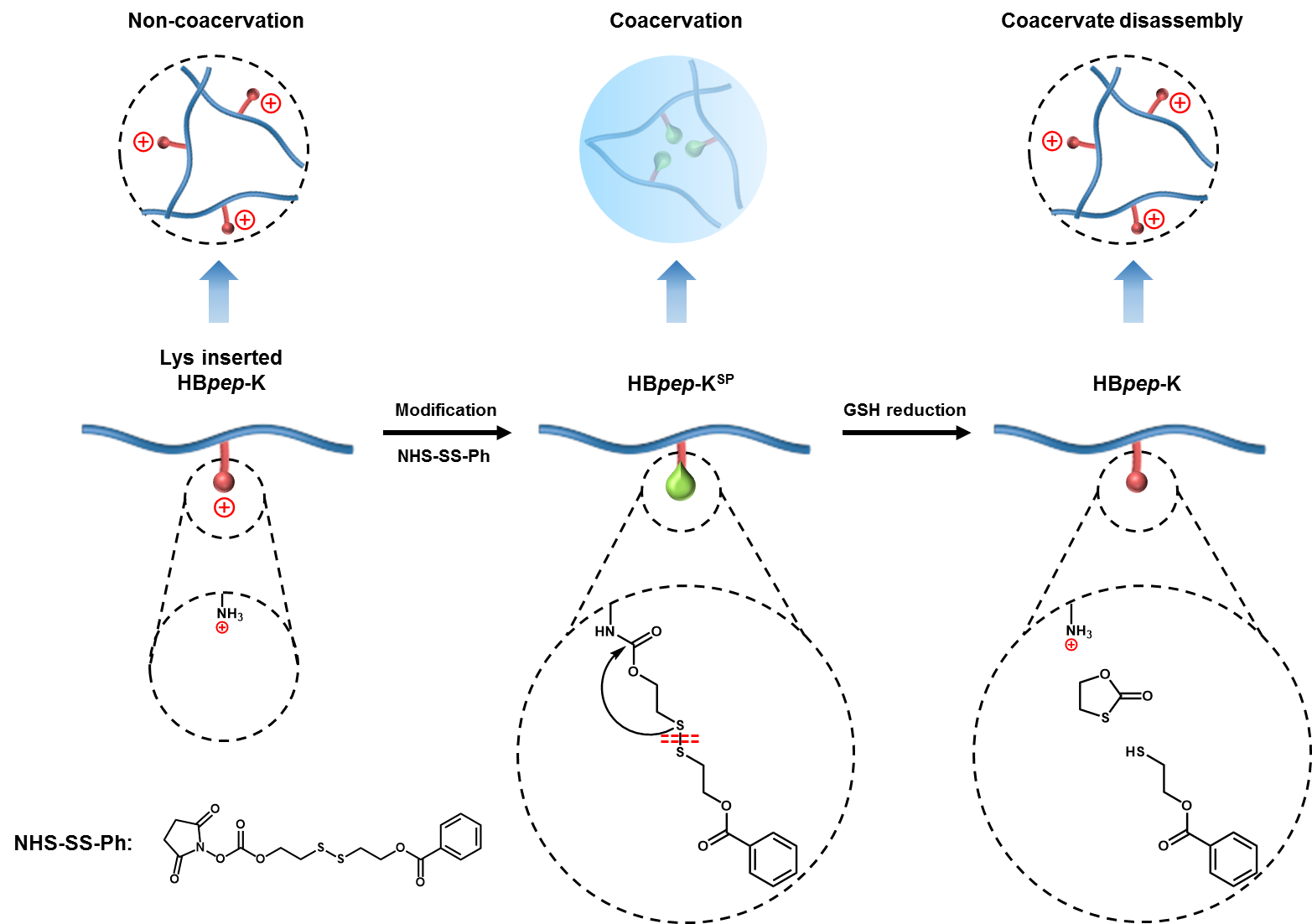

**Figure S1. Schematic illustration of self-immolative modification on the non-coacervating HB*pep*-K.** The modified HB*pep*-K^SP^ could be reduced by GSH and converted to non-coacervating HB*pep*-K upon reaching the cytosol, thereby triggering the coacervate disassembly and cargo release.

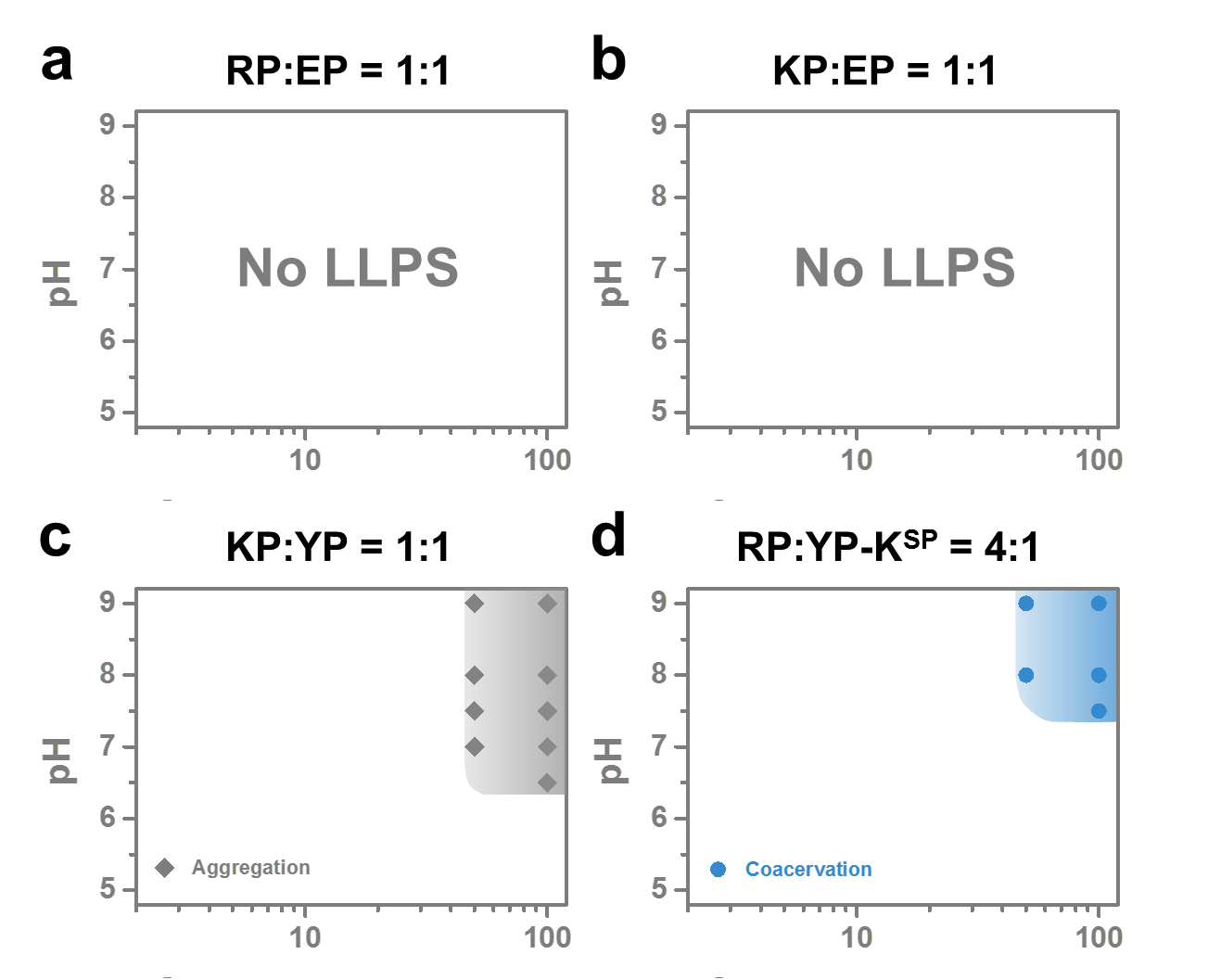

**Figure S2. Phase diagram of variant mixtures.** **(a)** RP:EP = 1:1, **(b)** KP:EP = 1:1, **(c)** KP:YP = 1:1, and **(d)** RP:YP-K^SP^ = 4:1 at the ionic strength (IS) of 100 mM, with the region of coacervation or aggregation shadowed in blue and grey, respectively.

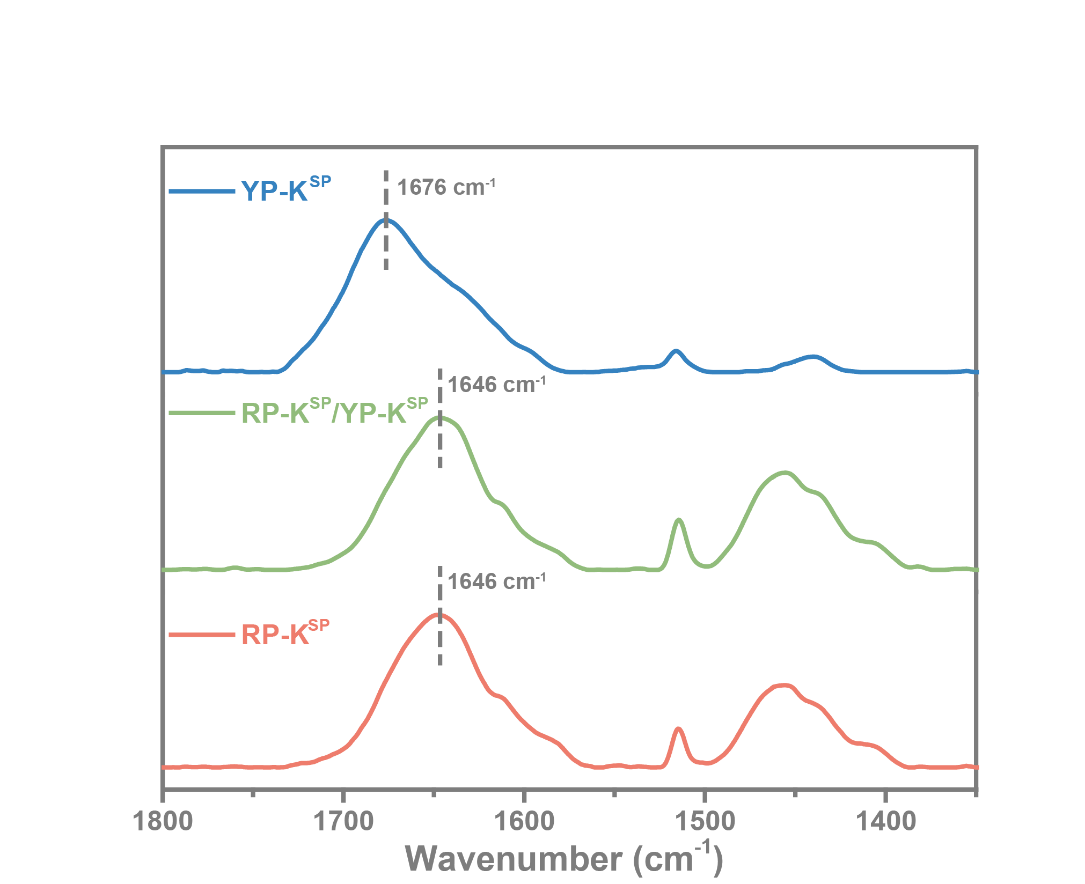

**Figure S3. ATR-FTIR of RP-K^SP^, RP-K^SP^:YP-K^SP^ = 1:1, and YP-K^SP^.** The amide I peak centered at 1676 cm^-1^ indicates β-turn structures, whereas 1646 cm^-1^ is attributed to disordered structures.

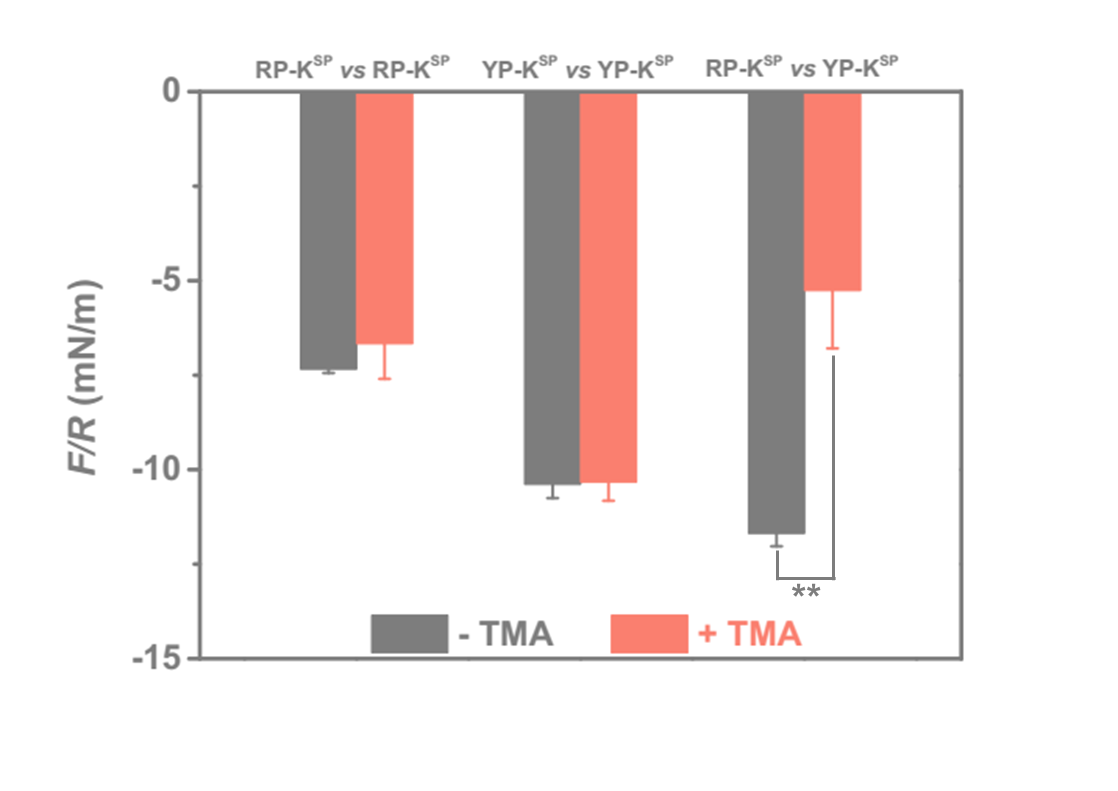

**Figure S4. Normalized adhesion force measured by SFA with RP-K^SP^ or YP-K^SP^ layers.** Measurements were conducted in buffer at pH = 7.0 and IS = 100 mM. Data are presented as the mean ± SD of *n* = 3 independent measurements; two-sided Student’s t-test, ***P*  <  0.01.

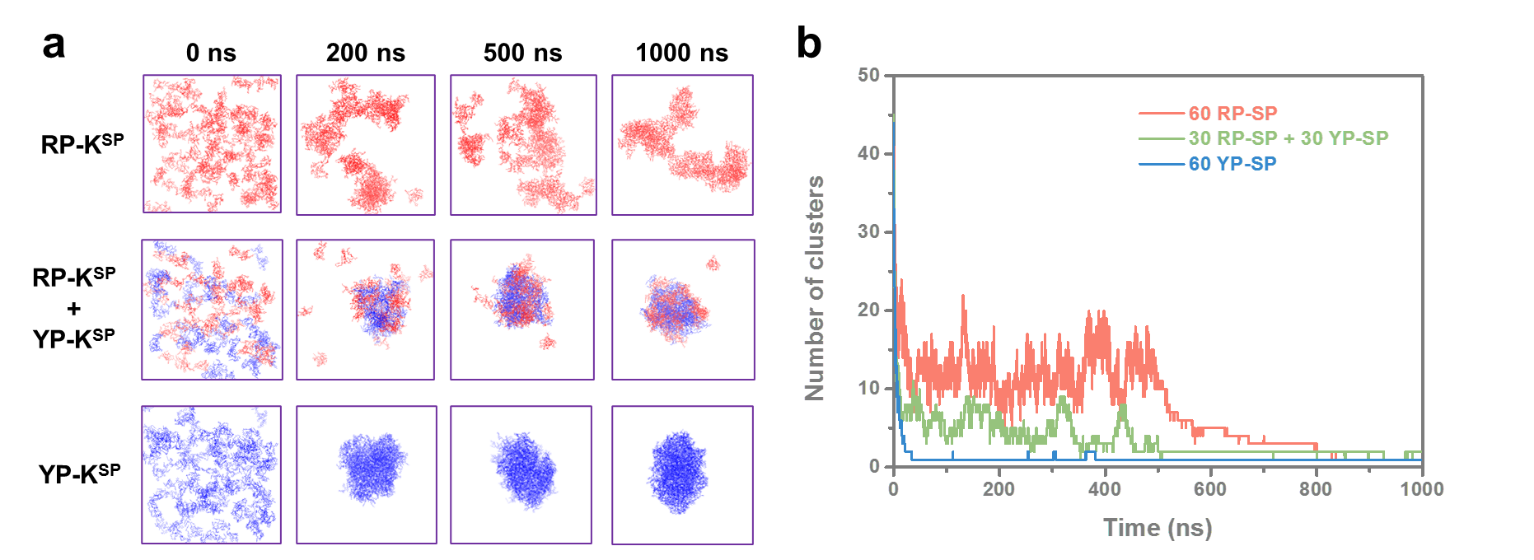

**Figure S5. Cubic MD simulations of RP-K^SP^, RP-K^SP^:YP-K^SP^ = 1:1, and YP-K^SP^. (a)** Snapshots of peptide molecules in three systems at various timepoints. **(b)** The number of clusters of three systems changes over time.

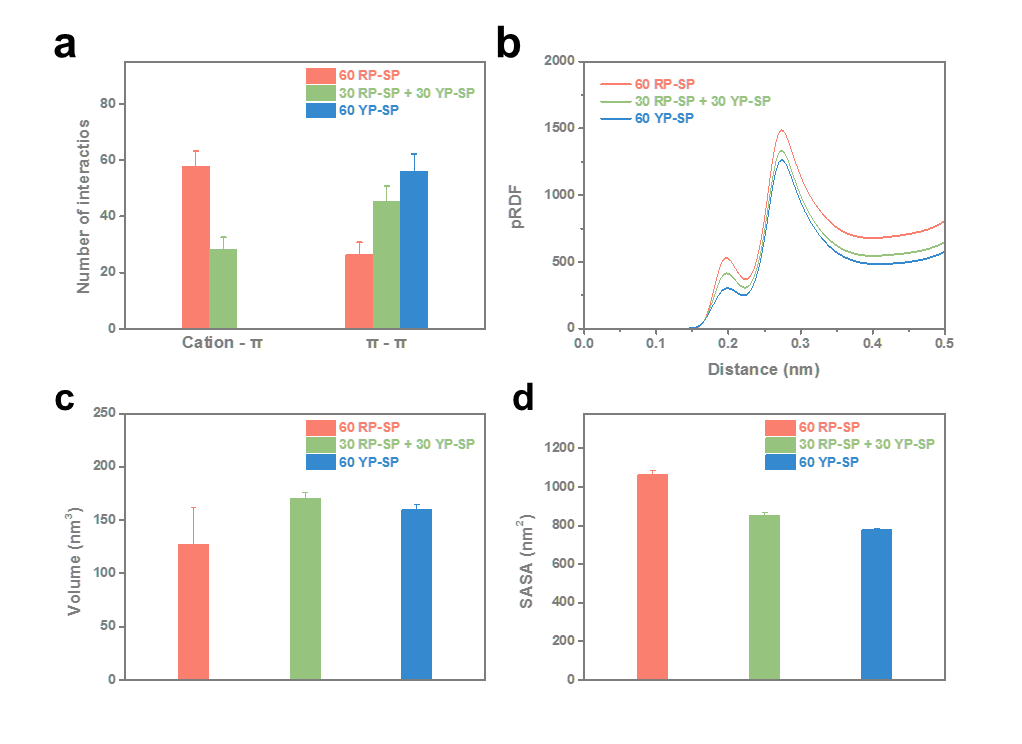

**Figure S6. Interactions and hydrations in the clusters formed by RP-K^SP^, RP-K^SP^:YP-K^SP^ = 1:1, and YP-K^SP^. (a)** The number of cation-π and π-π interactions in three clusters. **(b)** The proximal radial distribution functions (pRDF) of three clusters, suggesting the better hydration in RP-K^SP^, RP-K^SP^:YP-K^SP^ = 1:1 clusters compared to YP-K^SP^ **(c)** The volume of clusters formed by RP-K^SP^, RP-K^SP^:YP-K^SP^ = 1:1, and YP-K^SP^. caused by the differences in hydration. **(d)** The solvent accessible surface area (SASA) of three systems.

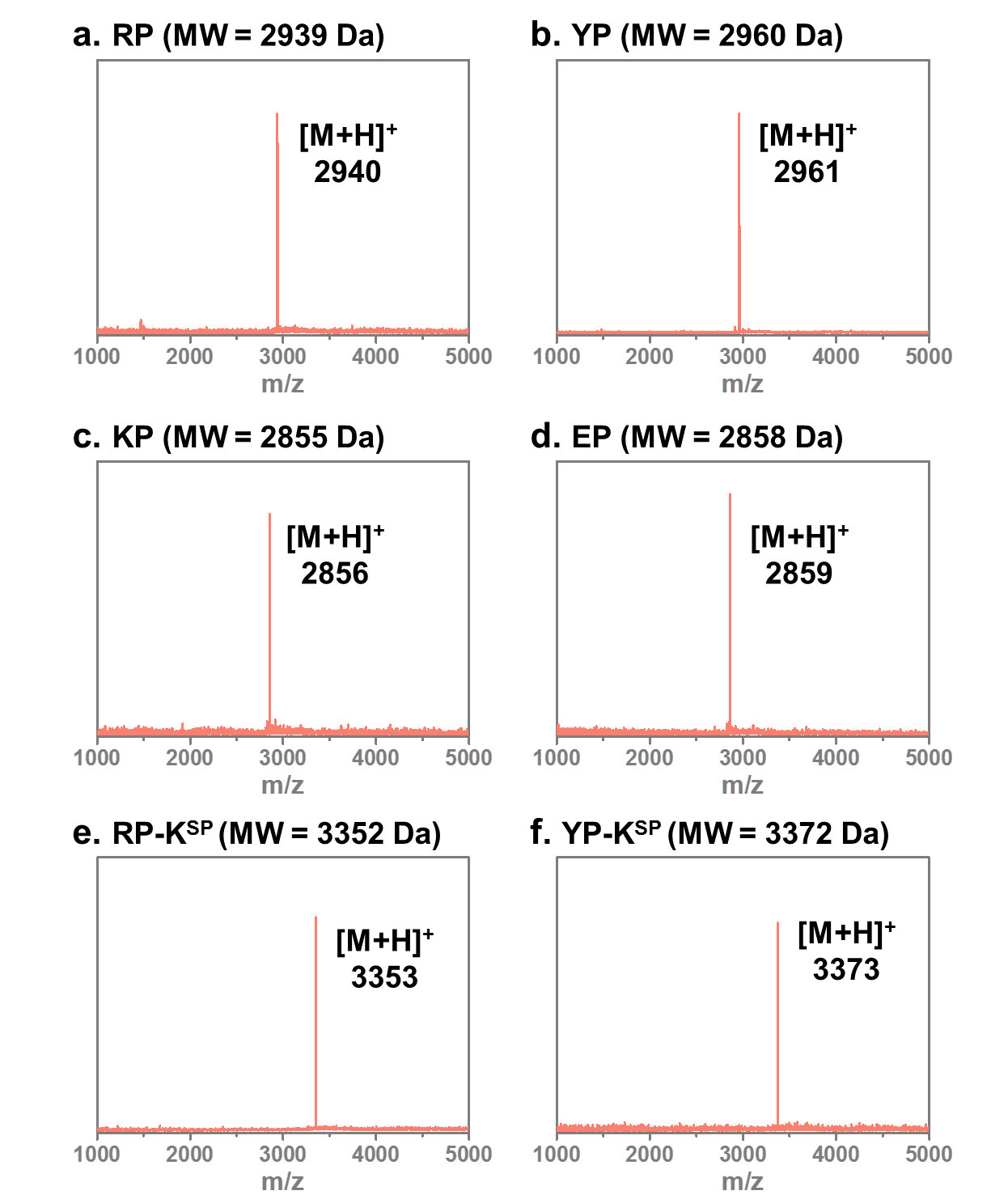

**Figure S7. MALDI-TOF spectra of HB*pep* variants with molecular weight (MW) indicated.** **(a)** RP, **(b)** YP, **(c)** KP, **(d)** EP, **(e)** RP-K^SP^, and **(f)** YP-K^SP^.

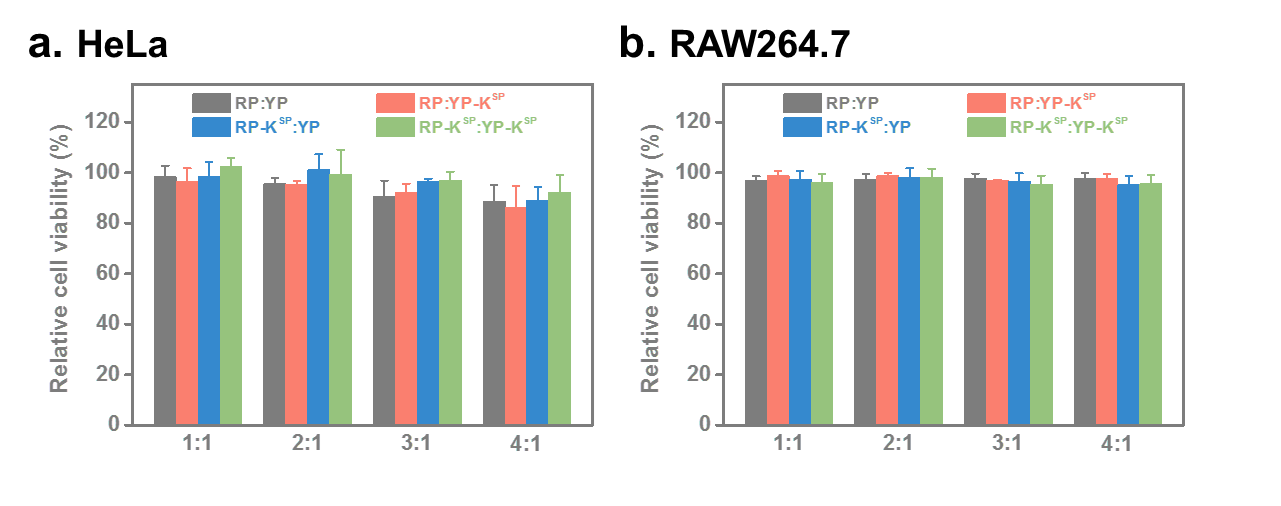

**Figure S8. Cytotoxicity of complex coacervates.** **(a)** Relative cell viability of HeLa cells treated with complex coacervates formed at various ratios. **(b)** Relative cell viability of RAW264.7 cells treated with complex coacervates formed at various ratios. Data are presented as the mean ± SD of n = 3 independent experiments.

**Supplementary Tables**

**Table S1.** Details of buffers used for LLPS study.

| **pH** | **Buffer salt** | **NaCl to fix 100 mM IS** |
| --- | --- | --- |
| 5 | 10 mM acetic acid | 94 mM |
| 6 | 10 mM sodium bicarbonate | 97 mM |
| 6.5 | 10 mM sodium dihydrogen phosphate | 84 mM |
| 7 | 10 mM sodium dihydrogen phosphate | 79 mM |
| 7.5 | 10 mM sodium dihydrogen phosphate | 74 mM |
| 8 | 10 mM sodium dihydrogen phosphate | 72 mM |
| 9 | 10 mM Tris | 99 mM |

**Table S2.** Details of each simulation system

| **No. of peptide** | **No. of water** | **No. of ions** | **No. of atoms** | **Time** | **Remarks** |
| --- | --- | --- | --- | --- | --- |
| 60 RP-K^SP^ | 164521 | 386 Na^+^, 566 Cl^-^ | 520795 | 2x1000ns | Cubic box |
| 30 RP-K^SP^ + 30 YP-K^SP^ | 164509 | 431 Na^+^, 521 Cl^-^ | 520489 | 2x1000ns | Cubic box |
| 60 YP-K^SP^ | 164529 | 476 Na^+^, 476 Cl^-^ | 520279 | 2x1000ns | Cubic box |
| 60 RP-K^SP^ | 56575 | 77 Na^+^, 257 Cl^-^ | 196339 | 2x1000ns | Slab box |
| 30 RP-K^SP^ + 30 YP-K^SP^ | 49051 | 87 Na^+^, 187 Cl^-^ | 173447 | 2x1000ns | Slab box |
| 60 YP-K^SP^ | 40163 | 117 Na^+^, 117 Cl^-^ | 146463 | 2x1000ns | Slab box |

**Table S3**. The sequence of sgRNA targeting SIRPα, and primers for amplifying SIRPα locus.

| **Name** | **Sequence** |
| --- | --- |
| sgRNA | CAAUGCUUGCAUAUUCUGUGGUUUUAGAGCUAGAAAUAGCAAGUUAAAAUAAGGCUAGUCCGUUAUCAACUUGAAAAAGUGGCACCGAGUCGGUGCUUUU |
| Forward primer | GCCAGTTGGTGGGTCAATA |
| Reverse primer | AGGCAGCTTTCTCCAGTTC |

**Supplementary** **References**

[1]	Y. Sun, S. Y. Lau, Z. W. Lim, S. C. Chang, F. Ghadessy, A. Partridge, A. Miserez, *Nat. Chem.* **2022**, *14*, 274-283.

[2]	Y. Sun, X. Xu, L. Chen, W. L. Chew, Y. Ping, A. Miserez, *ACS Nano* **2023**, *17*, 16597-16606.

[3]	S. H. Hiew, A. Miserez, *ACS Biomaterials Science & Engineering* **2017**, *3*, 680-693.

[4]	S. H. Hiew, H. Mohanram, L. Ning, J. Guo, A. Sánchez-Ferrer, X. Shi, K. Pervushin, Y. Mu, R. Mezzenga, A. Miserez, *Advanced Science* **2019**, *6*, 1901173.

[5]	J. Israelachvili, Y. Min, M. Akbulut, A. Alig, G. Carver, W. Greene, K. Kristiansen, E. Meyer, N. Pesika, K. Rosenberg, H. Zeng, *Reports on Progress in Physics* **2010**, *73*, 036601.

[6]	K. Deepankumar, Q. Guo, H. Mohanram, J. Lim, Y. Mu, K. Pervushin, J. Yu, A. Miserez, *Adv. Mater.* **2022**, *34*, 2103828.

[7]	X. Wu, Y. Sun, J. Yu, A. Miserez, *Communications Chemistry* **2024**, *7*, 5.

[8]	B. Lin, B. M. Pettitt, *The Journal of Chemical Physics* **2011**, *134*.

[9]	D. Sementa, D. Dave, R. S. Fisher, T. Wang, S. Elbaum-Garfinkle, R. V. Ulijn, *Angewandte Chemie International Edition* **2023**, *62*, e202311479.

[10]	G. L. Dignon, W. Zheng, Y. C. Kim, R. B. Best, J. Mittal, *PLOS Computational Biology* **2018**, *14*, e1005941.

[11]	F. J. Blas, L. G. MacDowell, E. de Miguel, G. Jackson, *The Journal of Chemical Physics* **2008**, *129*.

[12]	J. A. Maier, C. Martinez, K. Kasavajhala, L. Wickstrom, K. E. Hauser, C. Simmerling, *J. Chem. Theory Comput.* **2015**, *11*, 3696-3713.

[13]	W. L. Jorgensen, J. Chandrasekhar, J. D. Madura, R. W. Impey, M. L. Klein, *The Journal of Chemical Physics* **1983**, *79*, 926-935.

[14]	D. A. Case, H. M. Aktulga, K. Belfon, D. S. Cerutti, G. A. Cisneros, V. W. D. Cruzeiro, N. Forouzesh, T. J. Giese, A. W. Götz, H. Gohlke, S. Izadi, K. Kasavajhala, M. C. Kaymak, E. King, T. Kurtzman, T.-S. Lee, P. Li, J. Liu, T. Luchko, R. Luo, M. Manathunga, M. R. Machado, H. M. Nguyen, K. A. O’Hearn, A. V. Onufriev, F. Pan, S. Pantano, R. Qi, A. Rahnamoun, A. Risheh, S. Schott-Verdugo, A. Shajan, J. Swails, J. Wang, H. Wei, X. Wu, Y. Wu, S. Zhang, S. Zhao, Q. Zhu, T. E. Cheatham, III, D. R. Roe, A. Roitberg, C. Simmerling, D. M. York, M. C. Nagan, K. M. Merz, Jr., *J. Chem. Inf. Model.* **2023**, *63*, 6183-6191.

[15]	U. Essmann, L. Perera, M. L. Berkowitz, T. Darden, H. Lee, L. G. Pedersen, *The Journal of Chemical Physics* **1995**, *103*, 8577-8593.

[16]	M. J. Abraham, T. Murtola, R. Schulz, S. Páll, J. C. Smith, B. Hess, E. Lindahl, *SoftwareX* **2015**, *1-2*, 19-25.

[17]	G. A. Tribello, M. Bonomi, D. Branduardi, C. Camilloni, G. Bussi, *Comput. Phys. Commun.* **2014**, *185*, 604-613.

[18]	D. Y. Guschin, A. J. Waite, G. E. Katibah, J. C. Miller, M. C. Holmes, E. J. Rebar, in *Engineered Zinc Finger Proteins: Methods and Protocols* (Eds.: J. P. Mackay, D. J. Segal), Humana Press, Totowa, NJ, **2010**, pp. 247-256.

[19]	T. Wan, Q. Pan, Y. Ping, *Science Advances* **2021**, *7*, eabe2888.

[20]	L. Cai, X. Qin, Z. Xu, Y. Song, H. Jiang, Y. Wu, H. Ruan, J. Chen, *ACS Omega* **2019**, *4*, 12036-12042.
